# Cortical state contributions to neuronal response variability in the early visual cortex: A system identification approach

**DOI:** 10.1101/2024.09.17.613530

**Authors:** Jinani Sooriyaarachchi, Chang’an A. Zhan, Curtis L. Baker

## Abstract

Neurons in the early visual cortex respond selectively to multiple features of visual stimuli, but they respond inconsistently to repeated presentation of the same visual stimulus. Such trial-to-trial response variabilities are often treated as noise and addressed by simple trial-averaging to obtain the stimulus-driven response, though this approach is insufficient to fully remove the response variability. More importantly, response variability may primarily be caused by non-sensory factors, particularly by variations in cortical state.

Here we recorded and analyzed neuronal spiking activity in response to natural images from areas 17 and 18 of cats, along with local population neuronal signals, i.e. local field potentials (LFPs) and multi-unit activity (MUA). Single neurons showed highly varying degrees of trial-to-trial response variability, even when recorded simultaneously. We used a variability ratio (VR) measure to quantify the trial-wise differences in neural responses, and two cortical state indicative measures, a global fluctuation index (GFI) calculated using MUA, and a synchrony index (SI) calculated from LFP signals. We propose a compact convolutional neural network model with parallel pathways, to capture the stimulus-driven activity and the cortical state-driven response variabilities. The stimulus-driven pathway is comprised of a spatiotemporal filter, a parametric rectifier and a Gaussian map, and the cortical state-driven pathway contains temporal filters for MUA and LFPs. The model parameters are fit to best predict the spiking activity of each neuron.

The fitted model performed with a significantly higher accuracy in predicting neural responses compared to a basic model with a stimulus-driven pathway alone. The neurons with higher response variability benefited more from the cortical state-driven pathway compared to less variable neurons. These results show that different neurons may differ greatly in their variability and in the degree of their relationship to indicators of cortical state fluctuations.

**Author Summary:** Neuronal responses in the early visual cortex to repeated presentation of an identical stimulus can be highly variable across trials. The variable portion of these neuronal responses can in some cases be as large as the stimulus-driven response. The cortical state fluctuations that may underlie the response variabilities can vary continuously during a data recording session, and these dynamics are associated with population response signals such as local field potentials and multi-unit activity. Here we demonstrate that a model combining these cortical signals along with a visual stimulus processing pathway can predict single neurons’ responses significantly better than a model containing a stimulus-driven pathway alone. This improvement in predictive performance is heterogeneous across cortical neurons, and is much greater in neurons that exhibit greater trial-wise response variabilities. Overall, this work provides insights to understanding how visual cortex neurons not only respond to visual stimuli, but also interact with non-sensory events such as cortical state fluctuations.

## Introduction

A visual cortex neuron’s response depends on the characteristics of a visual stimulus in relation to its receptive field (RF). However, the neuronal responses elicited by repeated presentations of the same stimulus can be highly variable (Carandini, 2004; Tolhurst et al., 1983; Vogels et al., 1989). This trial-to-trial response variability is conventionally treated as noise and mitigated by simple averaging across trials. However, simultaneous recordings from multiple single neurons have demonstrated that such trial-wise response variabilities are correlated across neural populations (Averbeck et al., 2006; Zohary et al., 1994), suggesting that simple averaging places critical constraints on understanding biologically plausible visual information processing. In fact, cortical neurons can produce highly reliable spike trains for intracellularly injected current (Mainen & Sejnowski, 1995). Thus the observed response variability may not be solely caused by intrinsic noise of the neuron itself, but rather the activity of the network (e.g., cortical state, Harris & Thiele 2011) within which the neuron is embedded.

Recent attempts at developing models to explain this trial-to-trial variability in the visual cortex used population neural signals, e.g. local field potentials (LFP) and multi-unit activity (MUA), to infer cortical variability (Cui et al., 2016; Kelly et al., 2010). However these studies did not address the extent to which such response variability was heterogeneous across neurons, or the effects of cortical states in simultaneously recorded neurons. Understanding the nature of trial-to-trial response variability at the single neuron level could guide development of biologically plausible models to explain visual cortical computations and circuits. Furthermore, separating neurons’ responses to visual stimuli from brain state effects could facilitate improved characterization of neuronal sensory RFs.

The trial-to-trial variability of neural responses has been thought to arise from variability of synaptic inputs, reflecting the coordinated activities in the cortical network (Arieli et al., 1996; Carandini, 2004). These ongoing activity patterns were historically studied in terms of EEG, e.g. Berger’s observation of an 8-12Hz rhythm while his son sat passively with his eyes closed and a 14-30Hz rhythm when he opened his eyes (Berger, 1929). Such fluctuations of brain oscillations are often interpreted as reflecting cortical states, linked to multiple behavioral dynamics (Harris & Thiele, 2011). Although cortical states have been most studied as categorically distinct in relation to sleep stages (Brown et al., 2012) and depth of anesthesia (Liu et al., 2021; Ward-Flanagan et al., 2022), recent studies have indicated a more complex picture in which cortical state varies in a continuum during wakefulness (Reimer et al., 2014) due to factors including attention (Saalmann et al., 2007), working memory (Mendoza-Halliday et al., 2014), or general arousal (Constantinople & Bruno, 2011; Vinck et al., 2015). Recent studies on rodents demonstrated a relationship between brain-wide neural response variability and behavioral patterns including locomotor activity, pupil size and whisker movements (Musall et al., 2019; Stringer et al., 2019). Thus it seems likely that trial-to-trial variability may reflect the effects of cortical states, and their influences in sensory cortical function, rather than being just “noise”.

Disentangling the effects of cortical states on sensory information processing is challenging as the effects are confounded with the responses to the sensory stimuli. Moreover, brain state effects may not be uniform across a population of neurons. Some studies have demonstrated varying coupling of neuronal firing to the population activity, with more strongly coupled neurons showing greater activation during population-wide events such as top-down modulation (e.g. motor intention in saccade preparation) (Okun et al., 2015). Therefore it is important to study cortical state effects at the single neuron level to fully understand their differential effects.

Broadband responses from multielectrode recordings have been observed to co-vary with cortical states. These signals contain lower frequency components (0.5-100 Hz), commonly termed local field potentials (LFPs), which reflect synaptic inputs to a given cortical region (Buzsáki et al., 2012; Buzsáki & Draguhn, 2004), and more global EEG, which are thought to be indicative of brain states (Roohi-Azizi et al., 2017). The high frequency (300-5000 Hz) components (usually referred as to multi-unit activity, MUA) in nearby channels have also been found to predict single neurons’ spontaneous spiking activity (Okun et al., 2015; Schölvinck et al., 2015), suggesting that MUA may also be indicative of brain states. The LFP and MUA signals along with the visual stimulus information may be quantitatively analyzed jointly to uncover possible interactions between them (Cui et al, 2016; Kelly et al, 2010), rather than studying them in isolation.

Most early modelling efforts have assumed that cortical state acts either additively or multiplicatively on sensory neuronal responses. Multiplicative effects could conceivably result from top-down feedback related to attentional modulation (Armstrong & Moore, 2007), and some modeling efforts have assumed multiplicative (gain) modulation (Ecker et al., 2014). However, several studies modelled the cortical state modulatory effects on sensory processing as additive (Cui et al., 2016; Kelly et al., 2010; Murray, 2008; Thiele et al., 2009). In addition, one study revealed that a combination of both additive and multiplicative modulatory effects could better explain the trial-to-trial variability compared to either alone (Lin et al., 2015).

Previous studies were mainly aimed at improving accuracy in predicting neural responses but have seldom explored the effects of cortical states in estimating the RFs of the studied neurons. Furthermore, most of these studies used pre-defined LFP frequency bands like those conventionally used for EEG. However, the implied categorical nature of these frequency bands, and the assumed boundaries between them, might result in missing important relationships and information provided in the raw signals (Cohen, 2020).

Here we extended a system identification model of visual responses which estimates the receptive fields of early visual cortex neurons based on a simple convolution neural network (Nguyen et al., 2024). We incorporated the fluctuations of cortical states inferred from LFP and MUA signals recorded simultaneously from nearby electrode sites to explain the response variability of visual cortex neurons to repeated stimuli. The individual brain state effects on each neuron were captured using separate temporal filters applied to raw LFP and MUA signals in two parallel pathways.

We find that visual cortex neurons differ greatly from one another in their response variability, even when they are recorded simultaneously. Our complete system identification model including parallel pathways for visual stimuli, LFPs and MUA, can substantially improve the predictive accuracy of neuronal responses by better accounting for the response variability. Moreover, the improvement is greater for neurons with higher trial-to-trial response variability. Furthermore, we find that the quantitative improvement in predictive accuracy is also accompanied by improvements in the estimated RFs. The estimated LFP temporal filters extracted different important LFP frequencies uniquely for each neuron, showing differential effects of cortical states on different neurons including ones that were recorded simultaneously. These results thus demonstrate the significance of accounting for cortical state effects in characterizing the sensory responses at the individual neuron level.

## Materials and Methods

### Animal Preparation

Detailed descriptions of the procedures for animal surgery, recording, and maintenance have been published previously (St-Amand & Baker, 2023) and will only be summarized here. Adult cats of either sex were anesthetized with isofluorane/oxygen (3–5%), followed by intravenous (i.v.) cannulation. A bolus i.v. injection (5 mg·kg^−1^) followed by a continuous infusion (5.3 mg·kg^− 1^·h^−1^) of propofol was provided for surgical anesthesia. The animal was secured in a stereotaxic apparatus, and a craniotomy and small duratomy were performed over the cortical region of interest (Areas 17/18). The cortical surface was protected by 2% agarose and petroleum jelly. Body temperature was maintained using a thermister-controlled heating pad. The corneas were protected during the initial surgery using topical ophthalmic carboxymethylcellulose (1%), and subsequently, neutral contact lenses were inserted for long-term corneal protection. The eyes were brought into focus at the distance at which visual stimuli would be presented (57cm) using spectacle lenses, with artificial pupils (2.5mm) to further improve optical quality. Nictitating membranes were retracted, and pupils were dilated using topical phenylephrine hydrochloride (2.5%) and a mydiatic (atropine sulfate, 1%, or cyclopentolate, 1.0 %), which were re-applied daily.

The animal was connected to a ventilator supplying oxygen/nitrous oxide (70:30) after the completion of all surgical procedures. Gallamine triethiodide (i.v. bolus, to effect, followed by infusion, 10 mg·kg^−1^h^−1^) was used to induce and maintain paralysis. The propofol infusion rate was reduced (5.3 mg·kg^−1^h^−1^) and additional anaesthesia was provided with i.v. remifentanil (bolus injection, 1.25 μg·kg^−1^, followed by infusion, 3.7 μg·kg^−1^h^−1^). Vital signals (temperature, heart rate, expired CO2 and EEG activity) were monitored and kept at normal levels during recording experiments conducted over a period of 3-4 days.

All animal procedures were approved by the McGill University Animal Care Committee and are in accordance with the guidelines of the Canadian Council on Animal Care.

### Electrophysiology and Visual Stimuli

Extracellular recordings were obtained from 32-channel multielectrode probes (NeuroNexus, linear array: A1×32-6mm-100-177, linear edge: A1×32-Edge-5mm-100-177, or polytrode: A1×32-Poly2-5mm-50-177), using a Plexon Recorder (f_s_=31.25kHz) or OpenEphys (f_s_=30kHz) data acquisition system. Spiking responses (300Hz-5kHz) from a channel of interest were monitored “on-line”. All 32 channels of broadband neural data, as well as CRT photodiode signals (see below), were streamed to a hard disk for later off-line spike sorting using Kilosort 2.0 software (Pachitariu et al., 2016), MUA and LFP signal extraction, and temporal registration between stimuli and neural signals. The sorted neural spikes, MUA and LFP signals were then analyzed in detail as described below.

Probes were inserted in approximately vertical, near-columnar penetrations, so that the recorded neurons generally had similar receptive field location, preferred spatial frequency, and preferred orientation (DeAngelis et al., 1999; Hubel & Wiesel, 1962; Tootell et al., 1981). An initial estimate of the neurons’ receptive field location and dominant eye were identified using an interactively controlled bar stimulus. The non-dominant eye was occluded, and the CRT monitor centered approximately on the receptive field.

Visual stimuli were generated on a CRT monitor (NEC FP1350, 20”, 640 x 480 pixels, 150 Hz, 36 cd/m^2^) using custom software written in MATLAB (MathWorks, USA) with Psychophysics Toolbox (Brainard, 1997; Kleiner et al., 2007; Pelli, 1997) running on a Macintosh computer (MacPro, 2.66 GHz Quad Core Intel Xeon, 6 GB, NVIDIA GeForce GT 120, MacOSX 10.6.8). The monitor’s gamma nonlinearity was measured with a photometer (United Detector Technology) and corrected using inverse lookup tables. Stimulus onset/offset timing and electrophysiological recordings were temporally registered using a photodiode (TAOS, TSL12S) in a corner of the monitor screen. Photodiode signals were also used to verify the absence of dropped frames.

Detailed descriptions of the production of visual stimuli have been published (Talebi & Baker, 2012) and will only be summarized here. Natural image stimuli were constructed from monochrome 480×480 pixel subsets of photographs from the McGill Calibrated Color Image Database (Olmos & Kingdom, 2004). Images having low pixel standard deviations were discarded, since these images were nearly blank. We subtracted the mean luminance in remaining images and clipped to 8 bits. These natural images were the same as used previously (Nguyen et al., 2024; Talebi & Baker, 2012), except that in some experiments the image stimuli had higher RMS contrast, with the RMS contrast normalized or not. The observed final results were independent of these variations in the stimuli (S1 Fig) and will be considered jointly. Ensembles of 375 such images were presented in 5 sec trials. The CRT refresh rate was 150 Hz, with each stimulus frame presented twice, for an effective rate of 75 Hz. Stimulus image ensembles were divided into 3 sets, designed to be used for training, regularization, and testing (see below). Data collection was carried out in 5 trial blocks (henceforth referred to as “sweeps”) each consisting of 20 training ensembles, and 5 ensembles each for regularization and test sets. Image ensemble sets were presented in pseudorandom order over a period of ca. 45 minutes.

### Neural Data Processing

Neural data was processed in MATLAB (MathWorks, USA). The recorded broadband neural signals (sampling rate f_s_=31.25kHz with Plexon or f_s_=30kHz with OpenEphys) were filtered using Matlab’s *designfilt* and *filtfilt* functions. Neural signals were first low-pass filtered (<100Hz: passband frequency: 100, stopband frequency: 120, design method: ‘*kaiserwin*’, impulse response: ‘*fir*’, passband ripple: 0.1, stop band attenuation: 60), and also notch filtered to remove mains noise (60 Hz: half power frequencies: 59 and 61, design method: ‘*butter*’, impulse response: ‘*iir*’) to obtain local field potentials (LFPs), and high-pass filtered (>600Hz: passband frequency: 600, stopband frequency: 550, design method: ‘*kaiserwin*’, impulse response: ‘*fir*’, passband ripple: 0.1, stop band attenuation: 60) to obtain multiunit spiking activity.

To identify the recording channels in grey matter, current source density (CSD) analysis was employed on LFP signals extracted from responses to drifting sinewave gratings at varying spatial frequencies or orientations, which had been collected for conventional tuning curve measurements. These LFPs were analysed across the channels using the *CSD plotter* software (Pettersen et al., 2006) to identify the grey matter boundaries (S2 Fig). Channels with a prominent current sink in the CSD profile were specified as the granular layer, and sources above and below this sink were identified as the supragranular and infragranular layers respectively. Further analyses were performed only for grey matter channels.

Spike waveforms were extracted and classified using Kilosort 2.0 software (Pachitariu et al., 2016) to obtain single neuron spike time series, and to associate each neuron to a principal recording channel (Kilosort parameters include *Th*: [10,4], *minFR*: 1, *spkTh*: -5, *AUCsplit*: 0.9). Spike waveform clusters identified from Kilosort were manually accepted if the spike time correlograms showed a clear minimum near zero, spike amplitudes showed a normal distribution unique from the multi-unit spike amplitude distribution, and the shape of the spike waveform showed a clear distinction from noise. The acceptance, rejection and merging of spike clusters were carried out manually in Phy-GUI which is integrated with Kilosort. Spike-sorted neurons having an average frequency <1.0 spikes/sec in response to natural image ensembles were considered to be unresponsive and discarded from further analysis.

### Trial-to-trial response variability

As illustrated in Fig 1, a neuron’s response to repeated presentation of a natural image movie typically varied across trials. For each neuron we estimated a measure of its trial-by-trial variability as follows. For each of the n=20 repetitions of the regularization set, the Pearson’s correlation (*R*) between the neuron’s response to that repetition and the average response for the (n-1) other repetitions of the same stimulus were calculated. 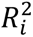 provided a reliability measure of each neuron’s response for a given trial *i* (*i=1, …, n*):

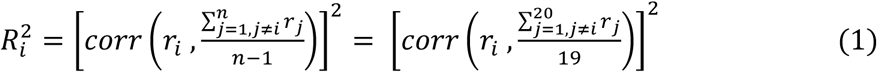

**Fig 1:**
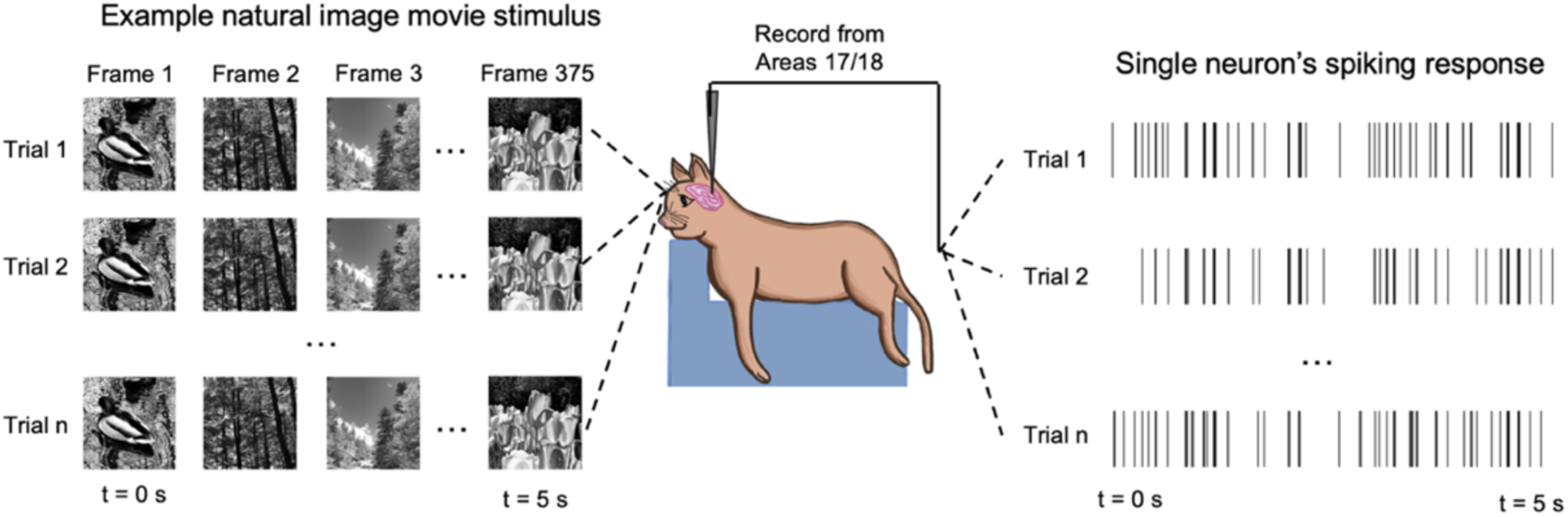
Trial-wise variability. Each natural image movie is presented across multiple (n) trials and the corresponding single neuron responses are compared to calculate a variability ratio, *VR* (Eq 1-3).

where *r_i_* refers to the neuron’s spiking response for the *i-*th trial, and *corr* denotes Pearson’s correlation. The average of these correlation coefficients was calculated:

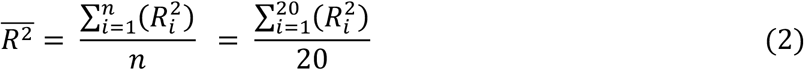

This 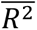 value is similar in principle to the “reliability ratio” of a neuron (Borst & Theunissen, 1999; Lesica et al., 2007). The variability ratio for each neuron was then quantified as:

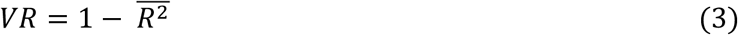

*VR* values could range from zero (perfect reproducibility) to unity (maximal trial-wise variability).

### Spike-LFP phase locking

We analysed the relationship of each neuron’s spiking activity to different LFP frequency bands, i.e. δ (0.5-4Hz), θ (4-7.5 Hz), α (7.5-12.5Hz), β (12.5-30Hz) and γ (30-100Hz) (Fig 2A and B). These LFP frequency bands were extracted in MATLAB by designing finite impulse response (FIR) bandpass filters using a Kaiser window applied to downsampled (x20) raw signals (i.e. for Plexon recorded data: from 31.25kHz downsampled to 1562.5Hz, OpenEphys recorded data: from 30kHz downsampled to 1500Hz). We then calculated a phase locking strength (*PLS_f_*) of each neuron’s firing to each filtered LFP band (Cui et al., 2016; Lachaux et al., 1999):

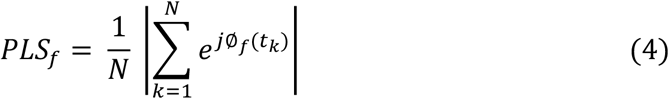

**Fig 2:**
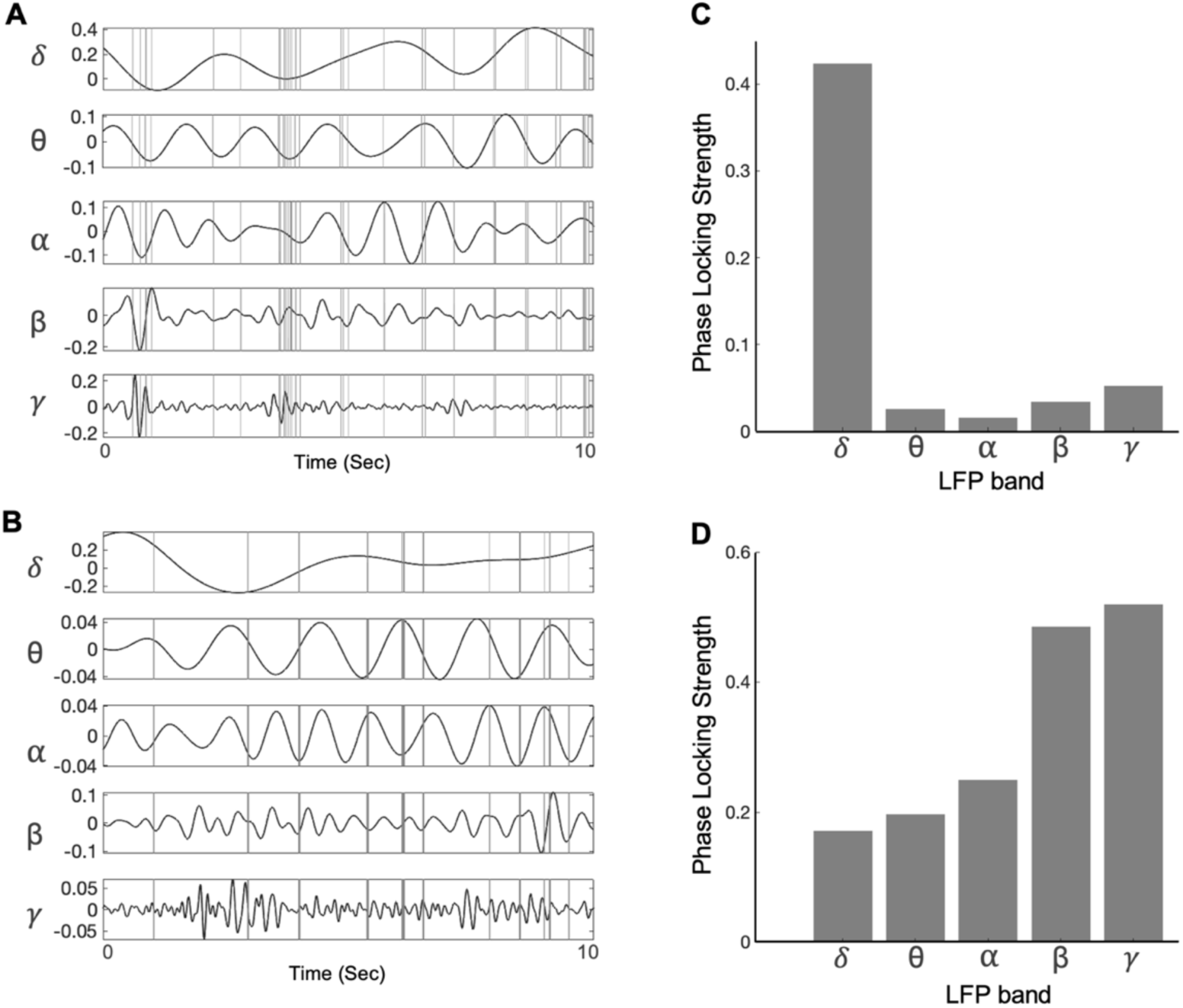
Spiking activity vs LFP frequency bands. **A)** Spiking activity of an example neuron 1 along with bandpass-filtered LFP signals recorded simultaneously. **B)** Similar to A, for example neuron 2. **C)** Phase locking strength (*PLS*) values (comparing spiking activity in relation to the phase of LFP bands) calculated for neuron 1. **D)** Similar to C, for example neuron 2.

where *f* indexes the LFP frequency band, *N* is the total number of spikes for the neuron, and ∅_f_(*t*_2_) is the phase of the LFP frequency band *f* at a given time *t_k_* of the *k*-th spike. ∅ _f_(t) was extracted using MATLAB’s *hilbert* function. PLS values could range from zero (no consistent phase relation) to unity (perfect phase locking).

### Cortical state dynamics

Local field potentials (LFPs) and multi-unit activity (MUA) show strong correlations with cortical state dynamics (Harris & Thiele, 2011). To identify the fluctuations of cortical state reflected by the population spiking activity across discrete time points, we extracted MUA (level crossings at ± 5 SD of the >600 Hz high-passed signal) from each of the channels in grey matter. To avoid confusion with responses to the visual stimuli, MUA-based cortical state analysis was performed during the 4 blank screen periods between the 5 sweeps. These MUA signals were binned (100 ms) to examine the up/down phases in network firing using a global fluctuation index (*GFI*, Eq 5) (Schölvinck et al., 2015). A comparatively higher GFI indicates prominent up/down phases characteristic of a synchronized cortical state. For each blank screen period, the binned firing rates were averaged across the channels to obtain channel-averaged firing rates (*FR*_34_678_). The *GFI* was taken as the ratio between the standard deviation and the mean ^of *FR*^34_678:

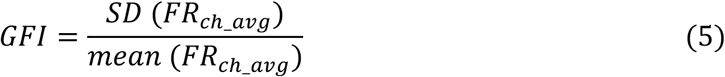

To analyse cortical state fluctuations in a continuous time scale throughout the data recording session, we calculated a synchrony index (*SI,* Eq 6) (Saleem et al., 2010) based on the LFP signals from the middle channel in layer IV identified from CSD analysis. The power spectral density (PSD) was first computed using Matlab’s *pspectrum* function (*type*: *’spectrogram’*, *FrequencyLimits*: [0 100], ‘*TimeResolution*’: 30 and ‘*OverlapPercent*’: 83.33) spanning throughout the data recording session. Then the Synchrony Index was calculated:

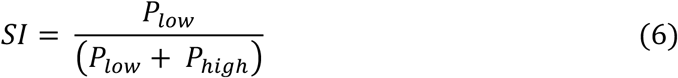

where *P_low_* refers to power in the frequency range of [0.5-15Hz] and *P_high_* refers to power in the frequency range of [15-100Hz] obtained from the PSD analysis. *SI* could vary from zero to unity, with higher *SI* values indicating a more synchronized brain state and lower *SI* values indicating a more desynchronized brain state.

### Model architecture and parameter estimation

We introduce a convolutional neural network model including parallel pathways: a stimulus-driven pathway to capture the neuron’s stimulus-dependent response (*R_stim_*), and cortical state pathways driven by two population signals, LFP and MUA, to capture the cortical state fluctuations (Fig 3).

**Fig 3:**
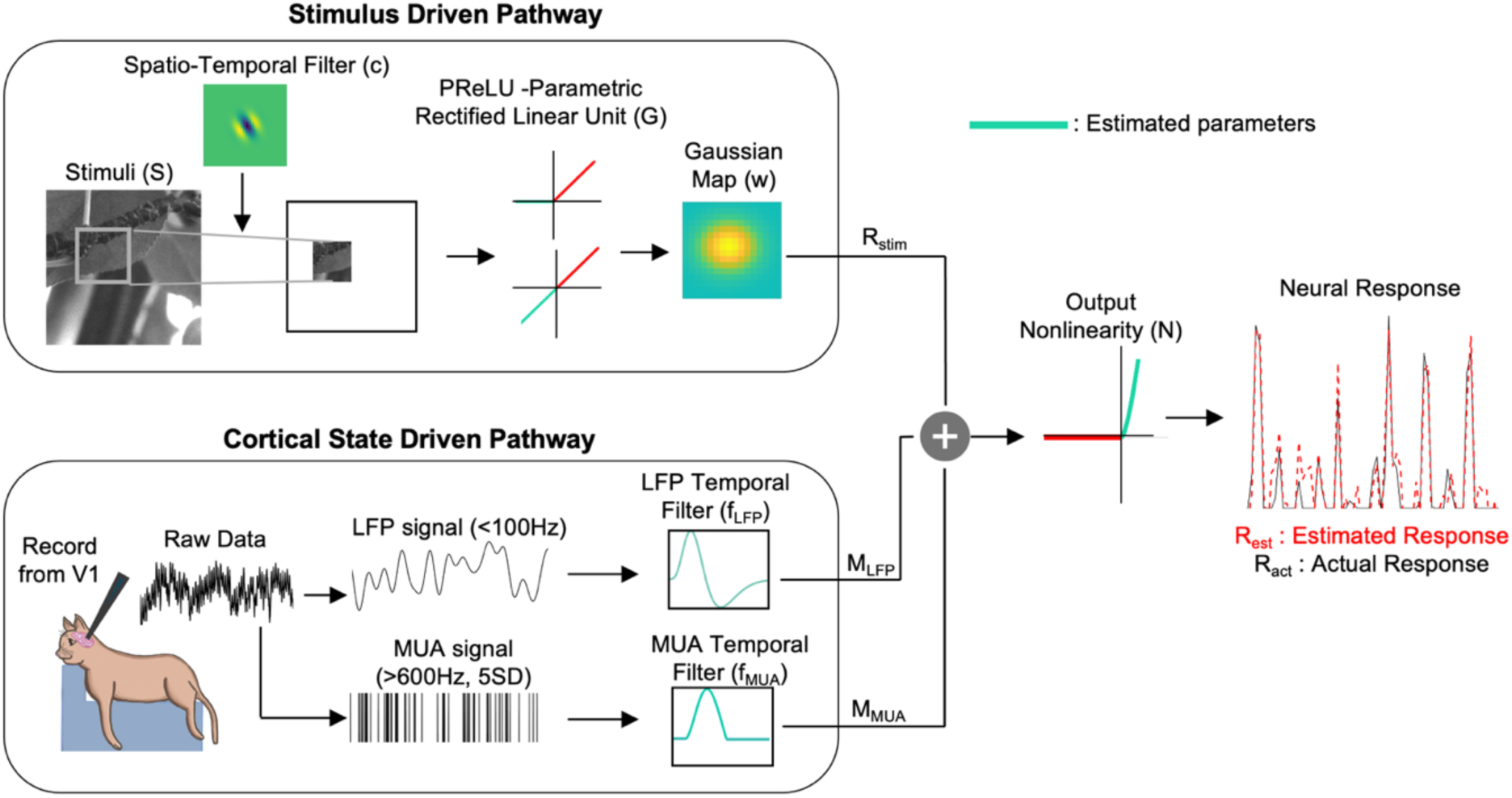
Model architecture for system identification. Visual stimuli (*S*) are convolved with a spatiotemporal filter (c), then passed through a PReLU (*G*) nonlinearity and a gaussian map (*w*) layer to provide the stimulus-driven response (*R_stim_*). Concurrent LFP and MUA signals provide inputs to the cortical state pathway, passing through LFP (*f_LFP_*) and MUA (*f_MUA_*) temporal filters to produce cortical state modulatory responses (*M_LFP_ and M_MUA_*). These signals summate with *R_stim_* and then are subjected to a final output nonlinearity (*N*) to produce the estimated neural response *R_est_*.

#### Stimulus-driven pathway

Details of the model architecture for the stimulus-driven pathway (Eq 7) have been described previously (Nguyen et al., 2024), so here will only be briefly summarized.

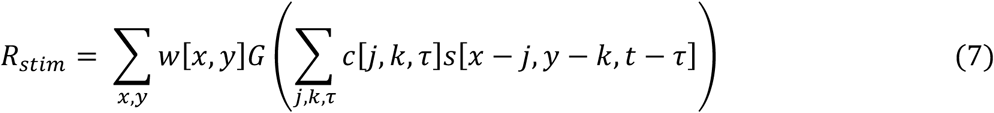

A cropped and downsampled stimulus *S* (see below) was first spatiotemporally convolved with the filter *c*. This was similar to the initial linear component of a linear-nonlinear model for a small region of the stimulus. The two-dimensional image output of this convolution, at a given time *t* was then passed through a static nonlinearity *G*, modelled using a Parametric Rectified Linear Unit (PReLU) with a single parameter PReLU_α_. The PReLU parameter, *α*, can take a value in a continuum from -1 to +1 indicating simple or complex type neural responses as well as intermediate types. The “map” layer *w* was modelled as a parameterized two-dimensional spatial Gaussian to identify the regions in visual space where the subunit filter *c* is operative.

#### LFP-driven pathway

Phase locking strength (PLS; Eq 4) values measured on a subset of neurons indicated that the spiking responses of different neurons could be phase-coupled to varying degrees with the components of different LFP frequency bands (Fig 2B and 2C). Therefore, we included LFP signals containing all frequencies up to 100Hz and let the model parameter-fitting procedure (below) estimate unique temporal filters applied to the LFP, for each neuron.

The LFP signals were first downsampled 10-fold (*fs_down_*= 3 kHz for OpenEphys and *fs_down_*= 3.125 kHz for Plexon-recorded datasets). These downsampled signals were then low pass filtered (<100 Hz), line noise was removed (notch filter at 60Hz) and they were segmented into windows of length 150 time points (corresponding to *T_window_* = 50 ms for OpenEphys and 48 ms for Plexon-recorded datasets). These LFP segments were computed for each frame of a natural image movie starting from the frame onset to a window length of 150 time points. Preliminary analysis with a subset of neurons indicated the sufficiency of using only 150 time points - longer window lengths did not improve the model prediction ability.

To ensure that the ability of the model to predict spiking activity from LFPs was not affected by any contamination from the spike waveforms, we avoided using LFPs from the same channel that the neuron was primarily recorded from (see Discussion). We evaluated the contribution of LFPs, gradually starting by incorporating LFPs from a nearby (located 100 μm away) channel to incorporating LFPs from all the channels (except the same channel) in grey matter. For recordings with polytrodes (having 2 columns of 16 recording sites), we used LFPs from the channels located along the same probe column as the recorded neuron, and we used every other channel (located 100μm apart) to be consistent with the linear array and edge recordings.

#### MUA-driven pathway

A neuron’s firing can be correlated to different degrees with nearby population firing, independent of stimulus conditions, which might occur due to non-sensory global variables including cortical state fluctuations (Okun et al., 2015). Therefore, we incorporated MUA from two nearby channels (100μm above and below) and included a MUA-driven cortical state pathway in our overall model. Unlike the LFP-driven pathway, here we did not use MUA signals from all the channels, because a given neuron’s firing correlates primarily with population firing that originates nearby (Cui et al, 2016). This idea was corroborated here by preliminary analyzes in a subset of neurons – the prediction accuracy of the system identification model did not further improve by including more MUA channels, and including all the channels resulted in estimated MUA temporal filter weights close to zero in channels further away from the primary channel (S3 Fig). The MUA from the same channel as the recorded neuron was always excluded to avoid potential signal contamination.

Signals from the two nearby channels were high pass filtered (>600Hz), and spikes exceeding a threshold of 5 standard deviations from the mean were detected to generate the MUA signal. This MUA signal was segmented into windows of 1500 time points (corresponding to *T_window_* = 500 ms for OpenEphys and 480 ms for Plexon-recorded datasets) starting at the onset of each natural image frame. These segments were downsampled 10-fold to yield segments of 150 data points in length (corresponding to *T_window_* = 50 ms for OpenEphys and 48 ms for Plexon-recorded datasets), similar to those from the LFP processing. An important issue is that in some cases, the spike waveform for one (sorted) neuron can appear in multiple channels. To avoid this kind of potential contamination, such “spillover” spikes were removed from the nearby channels before computing the MUA segments (S3 Fig).

#### Model architecture

LFPs and MUA, recorded simultaneously with spiking activity, both showed variations in *SI* and *GFI* (explained above) in our recorded data. We used these brain signals as indicators of cortical states to capture the variability in neural responses. To implement this, frames of natural image movies and the corresponding LFP and MUA segments were fed in parallel to the stimulus-driven pathway and LFP- and MUA-driven pathways of the model, respectively.

Overall activity from the cortical state-related signals was combined additively with responses from the stimulus-driven pathway, and subjected to a final output nonlinearity (*N):*

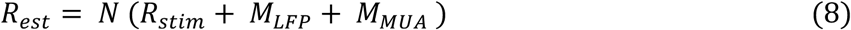

The output nonlinearity was modelled as a rectified power law, with gain (*g*) and exponent (*exp*):

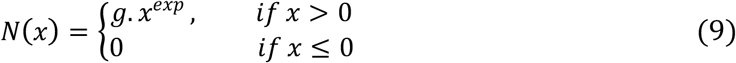

We also evaluated models in which the cortical state signals acted multiplicatively (Ecker et al., 2014; Goris et al., 2014), or with divisive normalization (Carandini & Heeger, 2011), but additive modulation of stimulus-driven neural activity by the cortical state-driven activity provided higher predictive accuracy (S4 Fig).

#### Model parameter estimation

Optimization of the model parameters was implemented in Python using tools from the TensorFlow 2.0.0 (Abadi et al., 2016) and Keras 1.1.2 (Chollet & others, 2018) packages. To avoid overfitting, we applied L2 regularization to the model coefficients from both LFP and MUA temporal filters and incorporated early stopping in the complete model with a patience value of 50 epochs. Preliminary analysis showed that the use of L2 regularization in the stimulus-driven pathway did not bring additional improvements in prediction accuracy. Model parameters were optimized as explained below.

First, a three-pass cropping procedure (Nguyen et al., 2024) was employed to identify regions of the stimulus that encompass the receptive field. We initially estimated the parameters of the model containing only the stimulus-driven pathway using full 480×480 stimulus images downsampled to 30×30 in the first pass. During the second pass, a square-shaped cropping window enclosing an area slightly larger than the apparent receptive field was identified and downsampled to 30×30 to re-train the model. In the final pass, we adjusted the cropping window based on the model estimated from the second pass to identify more accurate boundaries of the receptive field. We imposed a minimum cropping window size of 120 pixels to handle very small receptive fields. The final cropped stimulus from the third pass was downsampled to 30×30 to be used in further analyses as input to the stimulus-driven pathway.

The two parameters of the final output nonlinearity (Eq 9) were estimated in two stages, as previously described (Nguyen et al., 2024). In the first stage we set *N* to be a simple ReLU activation function and obtained a preliminary estimate of the receptive field filter and the filters for the LFP and MUA signals. For this optimization, we initialized the spatiotemporal receptive field filter with a Glorot normal initializer, PReLU *α* at 0.5 and the Gaussian map layer at the center of the image with standard deviation equal to the length of the image. For the second stage, we used the pre-trained receptive field weights from the first stage as initial conditions of the spatiotemporal receptive field (with other model parameters initialized similar to the first stage), and re-trained the model with *N* set to be an adaptive exponent output nonlinearity defined in a custom Keras layer with two trainable parameters, gain (*g*) and power law exponent (*exp*) as in Eq 9.

The models’ predictive accuracy for the hold-back test dataset was measured by the variance accounted for (*VAF*), computed as the square of the correlation between the neuron’s measured response (*R_act_*) and the response of the estimated model (*R_est_*) expressed as a percentage:

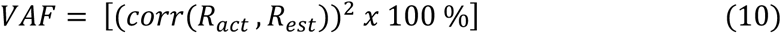

To quantify the trial-wise response variability explained by our cortical state-driven pathway we calculated a “*VAF_ongoing_*”. For this, we calculated the variable portion of the trial-wise response, which is considered as the “ongoing activity” (Schölvinck et al., 2015). The recorded ongoing activity (*Ongoing*_act_) for the test dataset was derived by taking the difference between each trial’s response and the trial-averaged response for 20 trials, which is considered as an estimate of the visual evoked response without brain state effects:

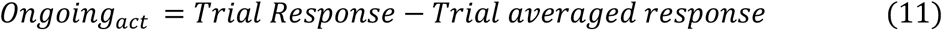

From the system identification model, we calculated the difference between the predicted responses from the full model containing the stimulus and cortical state-driven pathway and from the base model containing only the stimulus-driven pathway, which we term the ongoing activity estimated by the model (*Ongoing_est_*).

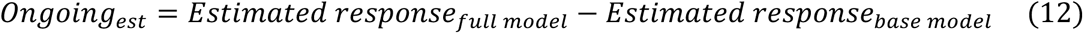

By comparing these two, we obtained the *VAF_ongoing_*:

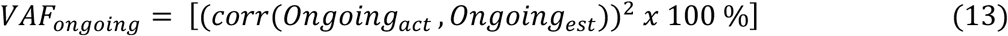

The *VAF_ongoing_* can vary between 0%, where the cortical state-driven pathway does not account for any of the observed ongoing activity, to 100%, where the cortical state-driven pathway completely accounts for the observed ongoing activity.

### Quality of receptive field estimates

We obtained receptive field restorations (henceforth termed “receptive fields”) by convolving the filter function (c) with the map function (w) (Nguyen et al., 2024). To quantify the statistical quality of estimated receptive fields, with or without incorporating brain state fluctuations, we calculated the z-score at the peak time lag of the receptive field using:

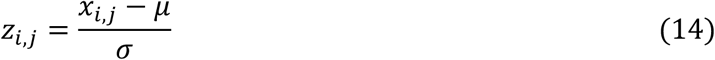

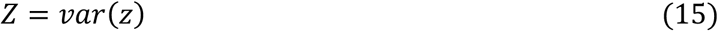

Here, *x_i,j_* is the *i,j-th* filter weight for the peak time lag of the receptive field, *μ* is the mean value of all the filter weights in the peak time lag, and *σ* is the standard deviation of the filter weights in the *zero*-th time lag of the spatiotemporal receptive field. Here *z* is a 2D array with values *z_i,j_* corresponding to each filter weight in the peak time lag. The variance of these values (Eq 15) was utilized in comparisons between models.

In the calculation above we used *σ* from the zeroth time lag as a measure of background noise of the estimated receptive fields (Eq 14). Since this lag is only nominally zero, but actually represents a time bin of 13.3 msec, it seemed prudent to check if there might be significant filter weights present at this lag depending on whether we incorporate brain state effects. To do this we confirmed the similarity of *σ* in the zeroth time lag of the estimated receptive fields before and after incorporating cortical state effects by using the Wilcoxon rank-sum test (S5 Fig).

### Dimensionality reduction of the filtered LFP power spectra

To examine the LFP frequencies of importance for different neurons, we further analysed the power spectra of recorded LFPs filtered by the estimated temporal filters for neurons which showed a VAF improvement >5% (S6 Fig). We then used a UMAP python package (McInnes et al., 2018) for nonlinear dimensionality reduction of the normalized power spectra (magnitude varying from 0 to 1), with parameters *n_neighbors=20, min_dist=0.1,* to obtain a high-dimensional graph. This graph was then used for the Louvain community detection algorithm (Blondel et al., 2008) to identify distinct clusters (*resolution = 1.0*). The *resolution* parameter was selected to maximize the modularity score which is a measure of the quality of clusters (S7 Fig). For visualization purposes, we projected the high-dimensional graph into two dimensions using UMAP. We compared the variability ratios and the VAF improvements of neurons in different clusters provided by Louvain clustering, to assess differential effects of different LFP power spectra.

## Results

We analyzed broadband neural data from 38 penetrations in Areas 17/18 of 14 anesthetized and paralyzed cats, which was recorded in multiple previous studies in the same lab (Gharat & Baker, 2017; Nguyen et al., 2024; St-Amand & Baker, 2023). The present study was based on analysis of 154 spike-sorted neurons (131 from OpenEphys and 23 from Plexon data acquisition systems) meeting the inclusion criteria (spike frequency >1.0 spikes/sec in response to natural image movies), along with simultaneously recorded LFP and MUA signals.

### Trial-to-trial response variability

We often observed single neuron spiking responses which differed across repeated presentations of the same stimulus. We quantified the extent of these trial-wise differences by a “variability ratio (*VR*)” (see Methods) which compares the similarity of individual trial responses to the trial-averaged response. Fig 4A shows spikes from an example neuron with a relatively low variability in response to repeated presentation of a 5 sec natural image movie. For this neuron the trial-averaged response and individual trial-wise responses were comparatively similar, yielding a low *VR* of 0.31. However, another example neuron as shown in Fig 4B exhibited much greater variability (*VR*=0.87).

**Fig 4:**
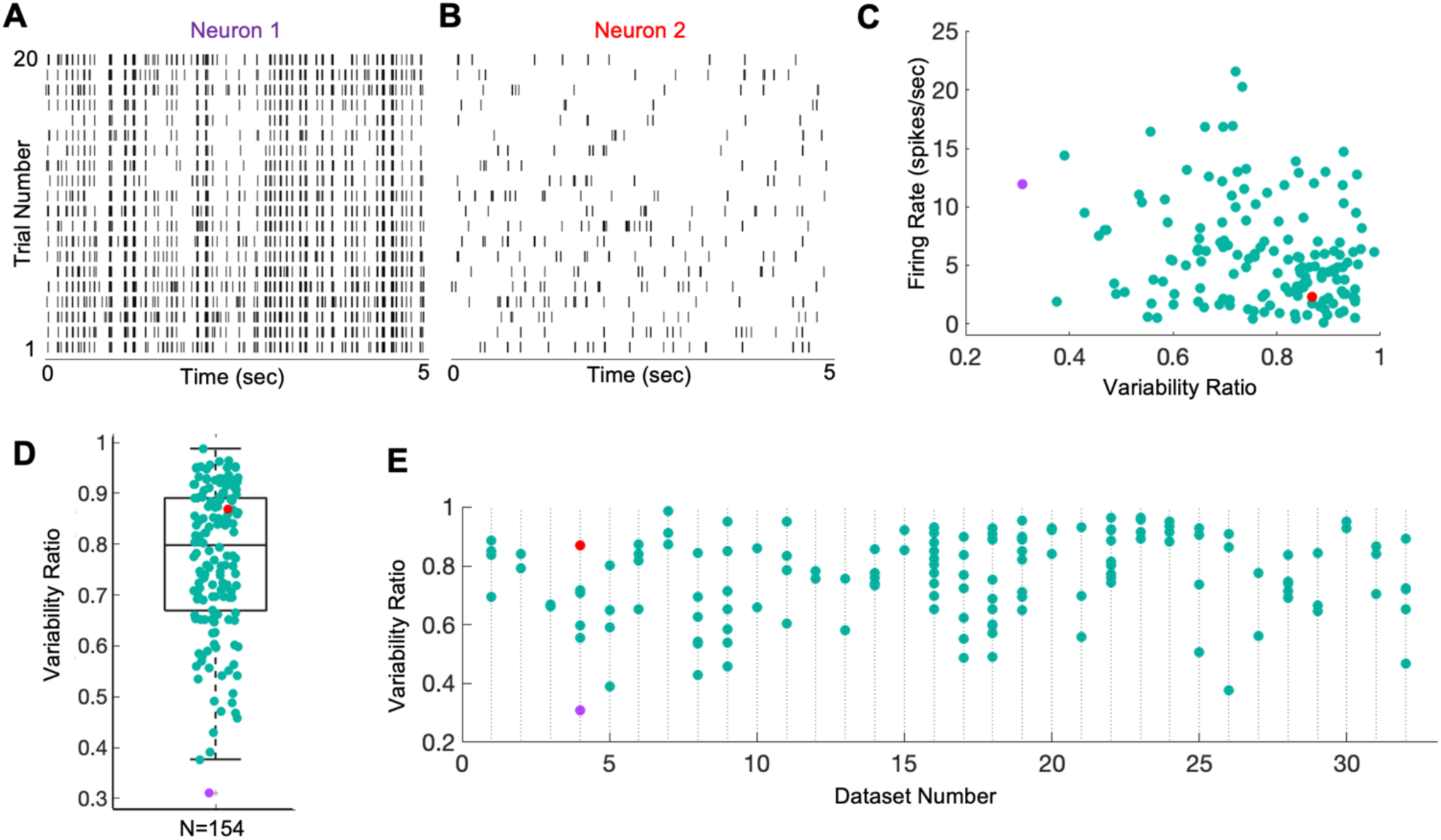
Trial-wise response variability of neurons. **A)** Spiking activity of a neuron with a low trial-to-trial variability (VR=0.31, the same neuron as in Fig 2A). **B)** Spiking activity of a neuron with a higher trial-to-trial variability (VR=0.87, same neuron as in Fig 2B) recorded simultaneously with the neuron in panel A. **C)** Scatter plot for the population of neurons (n=154) showing firing rate (FR) plotted against VR. Correlation coefficient between FR and VR is -0.21 (p=9.9e-03). **D)** Distribution of variability ratio across the population of neurons (n=154). **E)** Distribution of variability ratios of neurons across different datasets containing at least 2 simultaneously recorded neurons. In E, each data point on a column shows *VR* of a simultaneously recorded neuron in that dataset. In C, D, and E, data points from example neuron 1 is shown in purple and neuron 2 in red.

Comparing the two example neurons in Fig 4A and B, one might suppose that higher VRs and found in neurons with lower firing rates. To examine whether VRs are related to the mean FRs across the sample population, a scatter plot is presented in Fig 4C for the 154 neurons. While there is little evident relationship, statistical analysis shows a slight though significant negative correlation between the mean FRs and VRs (n=154, r= -0.18; p=0.03), suggesting that neurons with a low mean FR have a (slight) tendency to exhibit a higher VR and vice versa.

The *VR*s across the sampled population of neurons (n=154) are displayed in Fig 4D, showing *VR*s ranging from 0.31 to 0.98 with a mean of 0.73 (median 0.79) and standard deviation of 0.14 (inter-quartile 0.21). The high mean value of VRs suggest that the neural spiking activities are poorly predictable or explainable from the visual stimulus inputs alone. Traditional system identification models have usually ignored such variabilities and sought only to predict the trial-averaged firing rates (Nguyen et al., 2024).

It would seem conceivable that neurons with diverse VR values were those recorded at different times of higher or lower variability - in that case, one might expect that simultaneously recorded neurons would have similar VRs. Fig 4E shows a scatter bar plot of neuronal *VR*s for 32 datasets containing at least 2 simultaneously recorded neurons. In many instances, even the simultaneously recorded neurons showed very different amounts of trial-to-trial response variability, ranging from a maximum difference of 0.56 (dataset number 4 of Fig 4E) to a minimum of 0.01 (dataset number 3 of Fig 4E). This result shows that the sources causing the response variabilities can affect simultaneously recorded neurons differentially, and therefore neurons should be treated at an individual level to account for the observed response variabilities.

### Cortical state dynamics

The fluctuations of cortical network state might give rise to trial-to-trial response variability (Harris & Thiele, 2011). We thus analysed MUA and deep layer LFPs, as indicators of cortical network state fluctuations, using indices based on the firing patterns of simultaneously recorded populations of neurons and the LFP power in different frequency bands.

During a synchronized brain state, MUA shows prominent “up phases” where a majority of neurons are active and “down phases” where they are relatively silent (Destexhe et al., 2007; Harris & Thiele, 2011; Wilson, 2008). We analysed MUA signals during intersweep periods of spontaneous activity when the neuronal firing patterns are not affected by visual stimuli. LFP signals show high power in low frequency ranges and low power in high frequency rages during synchronized states and vice versa during desynchronized states (Harris & Thiele, 2011). Since LFP signals (<100Hz) are less affected by high frequency firing due to visual stimuli, we analysed LFPs throughout a recording session to continuously measure brain states, complementary to MUA that provided a measure of cortical states only during the intersweep intervals.

We computed MUA firing rate (100ms bins) for each of the channels in the grey matter during the four inter-sweep intervals. In some intervals (e.g. Figure 5A, intervals 1, 3, and 4), firing activity showed clear synchronization across channels, responding mainly during “up phases” and being completely silent during “down phases”. However, firing patterns in interval 2 were much less synchronized to the repeated stimuli, indicative of a relatively desynchronized state. The average firing rates across the channels (Fig 5B) confirmed the above impression that intervals 1, 3 and 4 show more pronounced fluctuations, with higher peaks for the “up phases”, than interval 2. The synchronized vs. desynchronized phases are further confirmed by quantifying a global fluctuation index, *GFI*, (Schölvinck et al., 2015), defined as the ratio of the standard deviation (SD) to the mean of channel-averaged firing rate (Methods, Eq 5), in Fig 5C. For additional examples, see also S8 Fig B, D, and F. Interval 2 showed a much lower GFI value than intervals 1, 3, and 4. Although these GFI values are calculated from responses when no visual stimuli were presented, previous studies show that the ongoing cortical state fluctuations can outlast visual stimulus presentations (Schölvinck et al., 2015).

**Fig 5.**
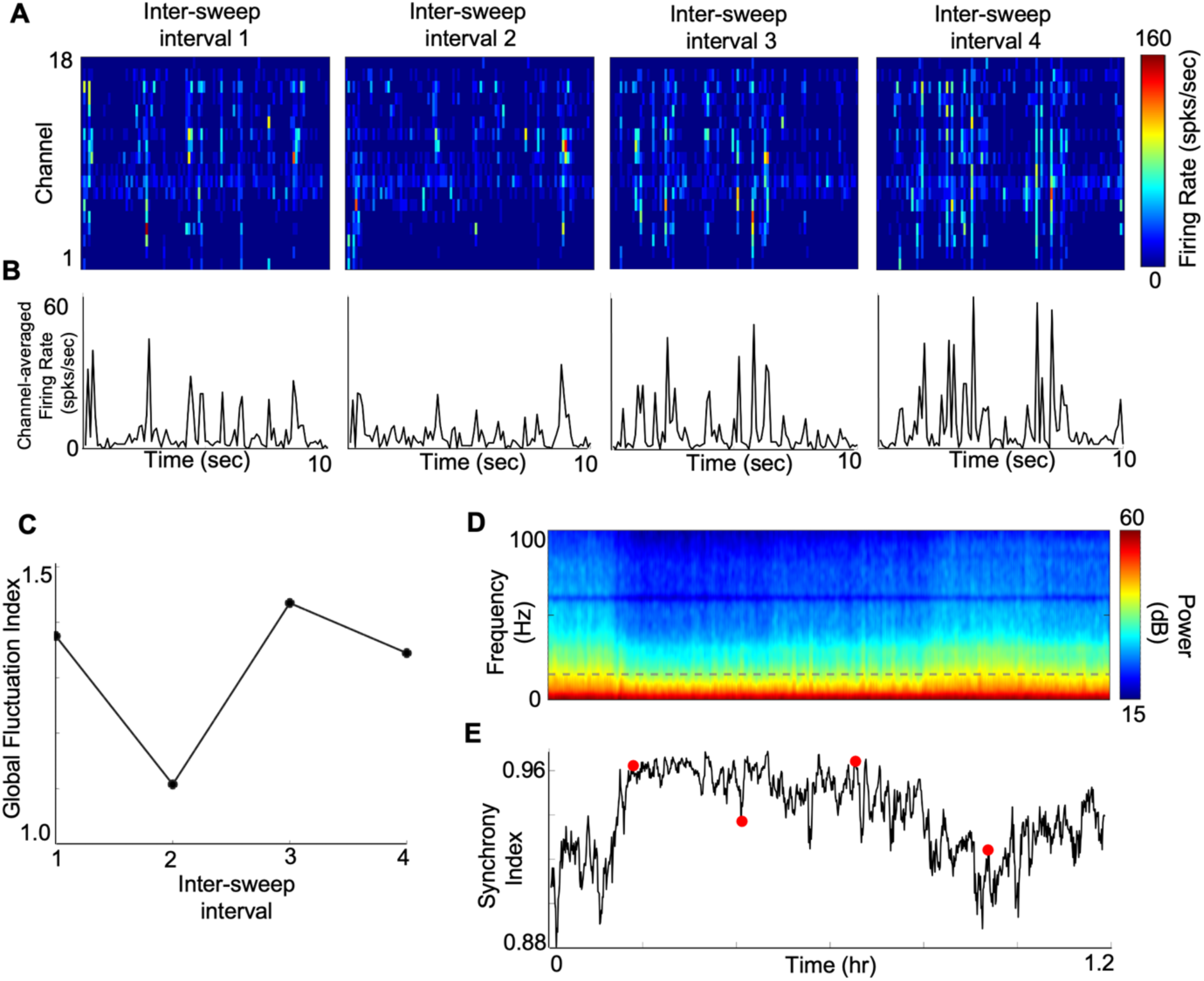
Cortical state fluctuations within data recordings. **A)** Binned firing rate of MUA across 18 channels in the grey matter (100 ms bins) during inter-sweep intervals while presenting a uniform grey screen. **B)** MUA firing rate averaged across channels in panel A. **C)** Global fluctuation index (GFI, Eq 5) using channel-averaged MUA firing rates in panel B. **D)** Power spectral density (PSD) of a deep layer LFP (middle channel of layer IV) across a data recording session (from same dataset as A-D). **E)** Synchrony index (SI, Eq 6) using PSD of panel D. Red data points indicate inter-sweep intervals.

Cortical state dynamics observed in terms of GFI could be obtained only at discrete times (i.e. during inter-sweep periods without any visual stimulation) during a data recording session. To identify network fluctuations in a more continuous manner, we analysed LFP signals spanning throughout a data recording session. Time frequency analysis of a deep layer LFP signal showed that the distribution of its power spectral density across frequencies varied greatly over time (Fig 5D, results from same recording session as for Fig 5A-C). We defined a synchrony index (*SI*) as the ratio of LFP power in low frequency ranges (0.5-15Hz) divided by the total power in low and high (0.5-100Hz) frequency ranges (Saleem et al., 2010). Overall, SI values in this example varied in a range approaching unity (range of 0.88 and 0.96) which could be due to the global cortical state under anesthesia. However, we observed continuous fluctuations of SI across time during the recording session (Fig 5E, also see S8 Fig for other examples) showing small rapid variations of network state dynamics which might result in the observed trial-to-trial spiking response variability. We observed a similar pattern of changes in the cortical state fluctuations from MUA and LFP when we overlaid them (Fig 5C, E) in all the data recordings. However, the statistical significance of correlation between the two measures varied in different data recordings (S8 Fig).

These fluctuations in GFI and SI show variations of the ongoing cortical state during a data recording session which might result in the observed neural response variabilities.

### System Identification Model

To examine how cortical state variations might influence neuronal activity, we incorporated candidate brain state indicators in a system identification model optimized to give best predictive performance of the neuronal spiking responses to visual stimuli. We developed the model (Fig 3) in stages incorporating successively more indicators of brain state.

#### Stimulus-driven (base) model

First, we consider the performance of the model for the stimulus-driven pathway alone (upper part of Fig. 3), without incorporating brain state effects, which we term the “base model”. With the recorded neural data and the corresponding stimulus input, the model parameters estimated for the example neuron 1 are presented in Fig 6A. The estimated model explained well the responses of this neuron (VAF 48.9%). The estimated filter weights contained a central inhibitory region surrounded by an excitatory region peaking at short time lags, which is reversed at larger time lags. The map layer, parameterized as a Gaussian, localized the activity of the filter. This example neuron had a PReLU alpha value of 0.62, corresponding to a nearly linear relationship indicative of a simple-type cell. The receptive field (i.e., the result of convolving the spatiotemporal filter weights with the Gaussian map) appeared less noisy and had a more clearly delineated excitatory surround. The convolutional output from the map layer passed through the final output nonlinearity (exponent 0.96) to generate the final neural response.

**Fig 6:**
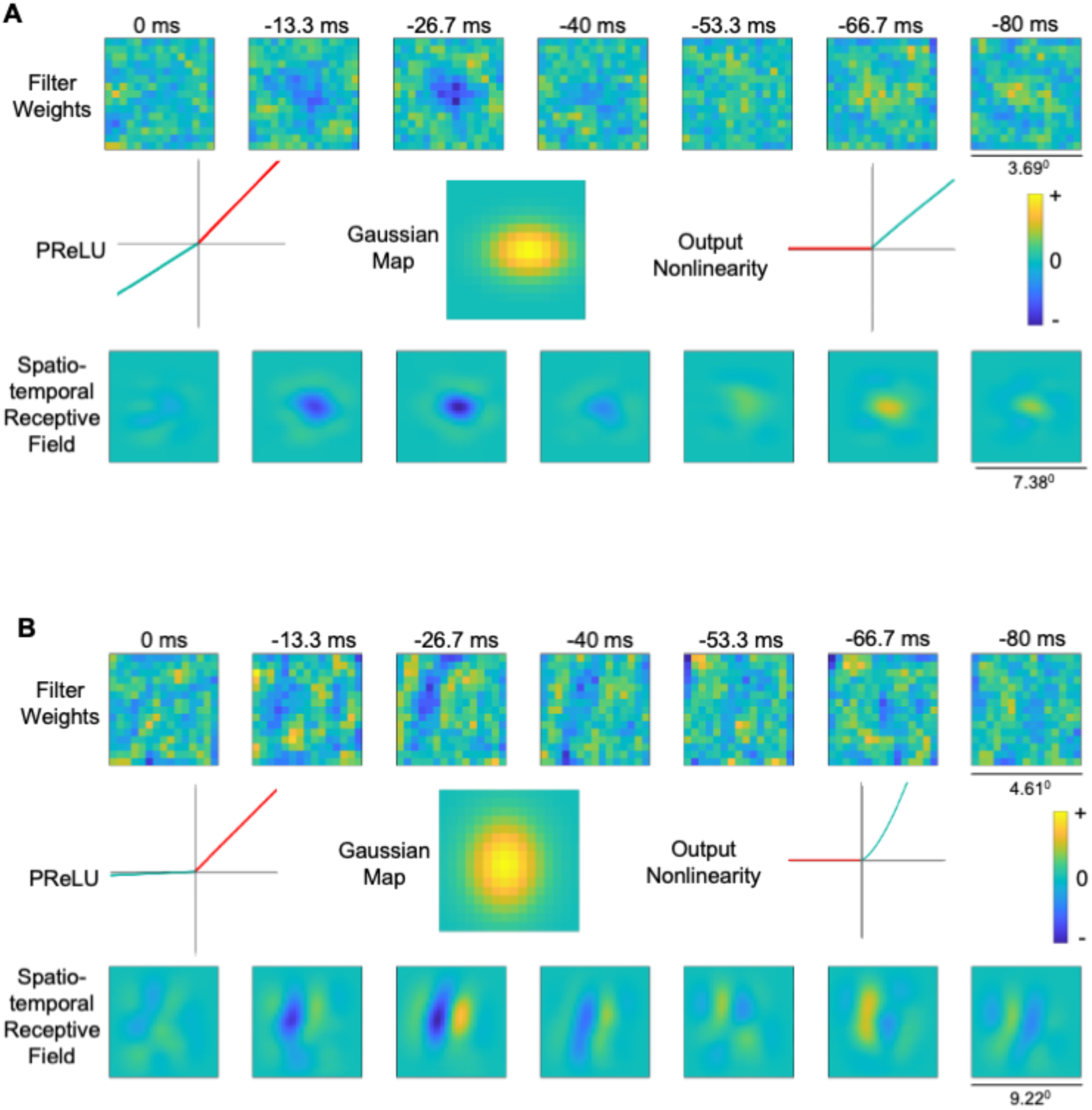
Estimated parameters for the stimulus-driven pathway for two example neurons. **A)** Estimated parameters from the stimulus-driven pathway for example neuron 1 (same neuron as in Fig 2B); spatiotemporal filter weights (top row), PReLU alpha, Gaussian map, Output nonlinearity and the final restoration of the receptive field (bottom row). **B)** Same as A, for example neuron 2 (same neuron as in Fig 2C)

The base model estimates for the second example neuron provided a VAF of 5.23%, much lower than for neuron 1. Fig 6B shows the spatiotemporal filter estimate (top row), PReLU alpha of 0.05 (indicative of a complex-like cell), estimated gaussian map, final output nonlinearity with exponent 1.50, and the receptive field (bottom row). The estimated receptive field for this neuron was spatially Gabor-like, i.e. with elongated, alternating excitatory and inhibitory regions, which reversed polarity at larger time lags.

Neurons with VAF<1% using the model consisting of stimulus-driven pathway only, were discarded from further analysis to help ensure reliability of the estimated model parameters.

We reasoned that the lower VAF for the second neuron could be a result of its higher trial-to-trial response variability (Fig 4A, B). Thus, we next incorporated the influence of cortical state indicators to see whether there are improvements in the model performance, i.e. predictive VAF accuracy.

#### Performance improvement with cortical states estimated using LFPs

Since cortical state fluctuation patterns can be detected with LFPs recorded simultaneously with spiking activity (Banerjee et al., 2012; Kelly et al., 2010), we incorporated LFP signals in a cortical state-driven pathway, parallel to the stimulus-driven pathway in our model architecture (Fig 3) to predict trial-to-trial response variability. For each of the neurons we re-trained the base model with the cortical state-driven pathway, and the optimization algorithm estimated a temporal filter to enhance LFP frequencies from a nearby channel related to the trial-wise variable response. Here we will first examine results in detail for the two example neurons, and then consider results for the sample population.

The estimated temporal filter for the first example neuron (Fig 7A) had a relatively low gain (± 0.001) with little evident temporal structure. However, the LFP temporal filter for the second neuron (Fig 7B) was ∼20 times larger in gain (± 0.02), with clear positive weighting of shorter time lags, and a negative peak around 25 ms. The filter weights for the first example neuron were relatively small for all the time lags and therefore the filter would dampen the LFP signals passing through it. For the second example neuron, the measured filter weights were relatively larger in magnitude, with an evident temporal structure, such that the filter could retain relevant frequencies for the neuron and dampen less important frequencies.

**Fig 7:**
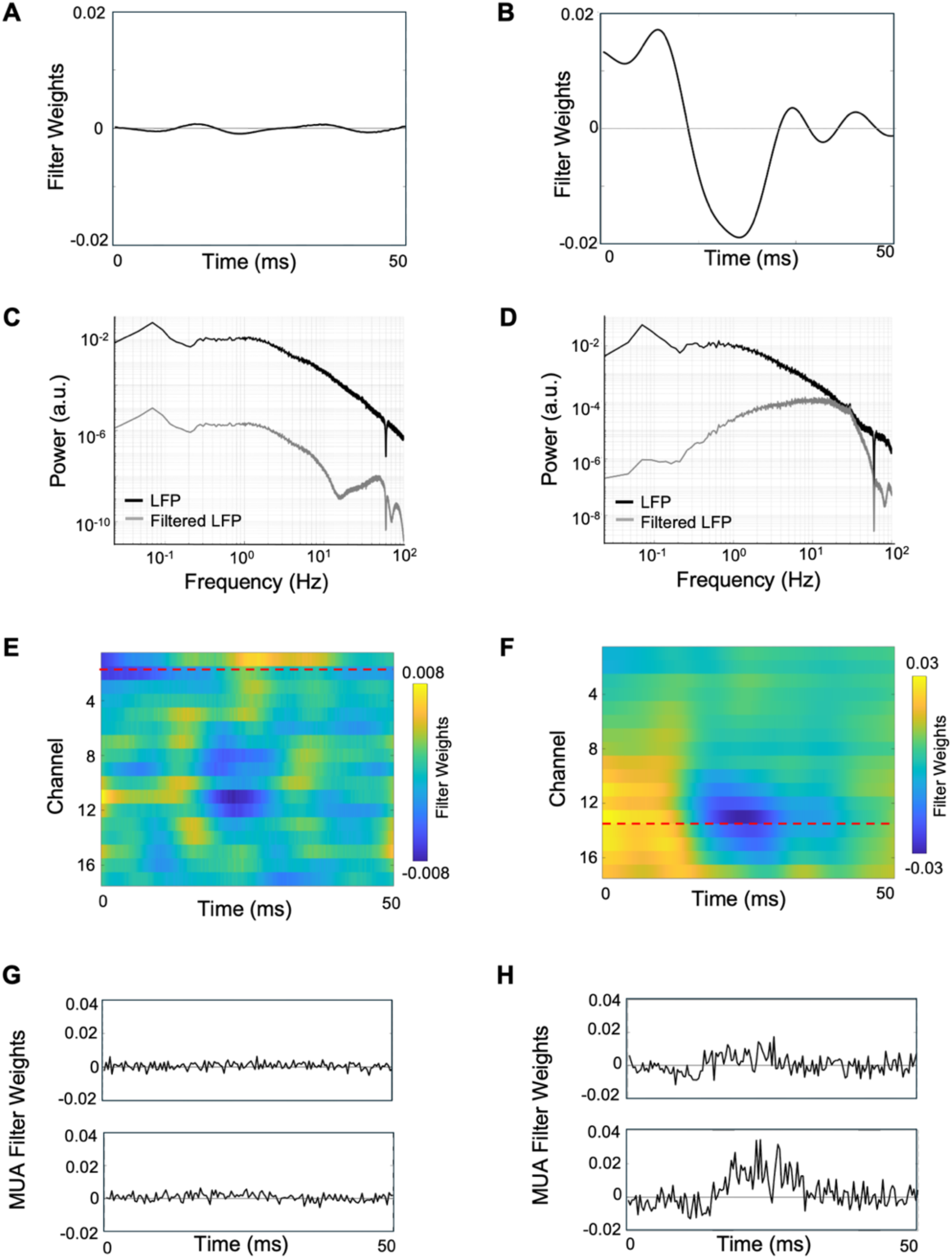
Estimated parameters for the cortical state-driven pathways, for two example neurons. **A)** Estimated temporal filter from LFP-driven pathway for example neuron 1 when using LFP only from the channel above. **B)** Similar to A, for example neuron 2. Note ordinate scales are same for A and B, to facilitate comparison. **C)** Frequency spectrum of LFP before (black) and after (grey) applying the temporal filter shown in panel A. **D)** Similar to C, for neuron 2. **E)** Estimated temporal filters from LFP-driven pathway for neuron 1 when using LFPs from all the grey matter channels. Dashed red lines indicate recording channel of the neuron. **F)** Similar to E for neuron 2. **G)** Estimated temporal filters from MUA-driven pathway for neuron 1 when using MUA from the above and below channels. **H)** Similar to G for neuron 2. Note ordinate scales are same for G and H. (See also S6, S9 Fig for additional examples).

To identify the relevant frequency bands of importance for each neuron we calculated the frequency spectrum of the LFP signal filtered through the estimated temporal filter. For the first example neuron, the filtered LFP had a similar power spectrum as the raw LFP, but with a broadband reduction in power (Fig 7C). The power spectrum for this filtered LFP had higher power in the delta frequency range (0.5-4Hz), comparable with the LFP-spike phase locking values for the neuron, showing an abundance of spikes phase-locked to the delta band (0.5-4Hz) (Fig 2C). For the second example neuron, the filtered LFP spectrum’s shape was markedly different compared to the raw LFP power spectrum, with greater power in the higher frequency ranges including beta (15-30Hz) and gamma (30-100Hz) compared to the lower frequencies (Fig 7D). This behavior was comparable with the LFP-spike phase-locking values for the neuron, showing spikes more phase-locked to the higher frequencies (Fig 2D). Overall, phase-locking strengths for the second example neuron had relatively larger values (0-0.6), and showed higher spike-LFP coherence (Fig 2B, C), compared to the first neuron (0-0.4),

Consistent with these observations, the predictive VAF accuracy for the second example neuron improved from 5.23% to 8.14% when the LFP pathway was included in the model. This improvement was higher than for the first neuron, which only improved from 48.9% to 49.1%. These observations suggested that the second neuron shows higher cortical state effects compared to the first neuron, as evident from its larger trial-to-trial variability ratio and higher VAF improvement with the incorporation of LFP signals in the system identification model.

To further examine how neurons varied in terms of the LFP frequencies filtered by the estimated temporal filters, and whether this was related to the VAF improvements, we examined the power spectra of filtered LFPs of all the sampled neurons which showed VAF improvements >5%. We excluded neurons with smaller VAF improvements since their temporal filter values were close to zero (e.g. Fig 7A) - the filtered LFP spectra for a representative subset of the remaining n=61 neurons are shown in S6 Fig. Non-linear dimensionality reduction of the filtered LFP power spectra (see Methods) identified three distinct clusters (Fig 8A). One cluster showed a low-pass filtered LFP power spectrum (“class 1“: n=22, Fig 8B, purple), while a second group showed a broader band-pass filtered spectrum (“class 2“: n=15, Fig 8C, yellow) and a third cluster showed a narrow band-pass filtered spectrum (“class 3“: n=24, Fig 8D, cyan). However, the VAF improvements and the variability ratios of the neurons in these three classes of filtered LFP power spectra were not significantly different (Fig 8E-F, p>0.05). These observations suggest that different neurons can show a range of response variabilities and VAF improvements independent from the LFP frequency ranges that influence them.

**Fig 8:**
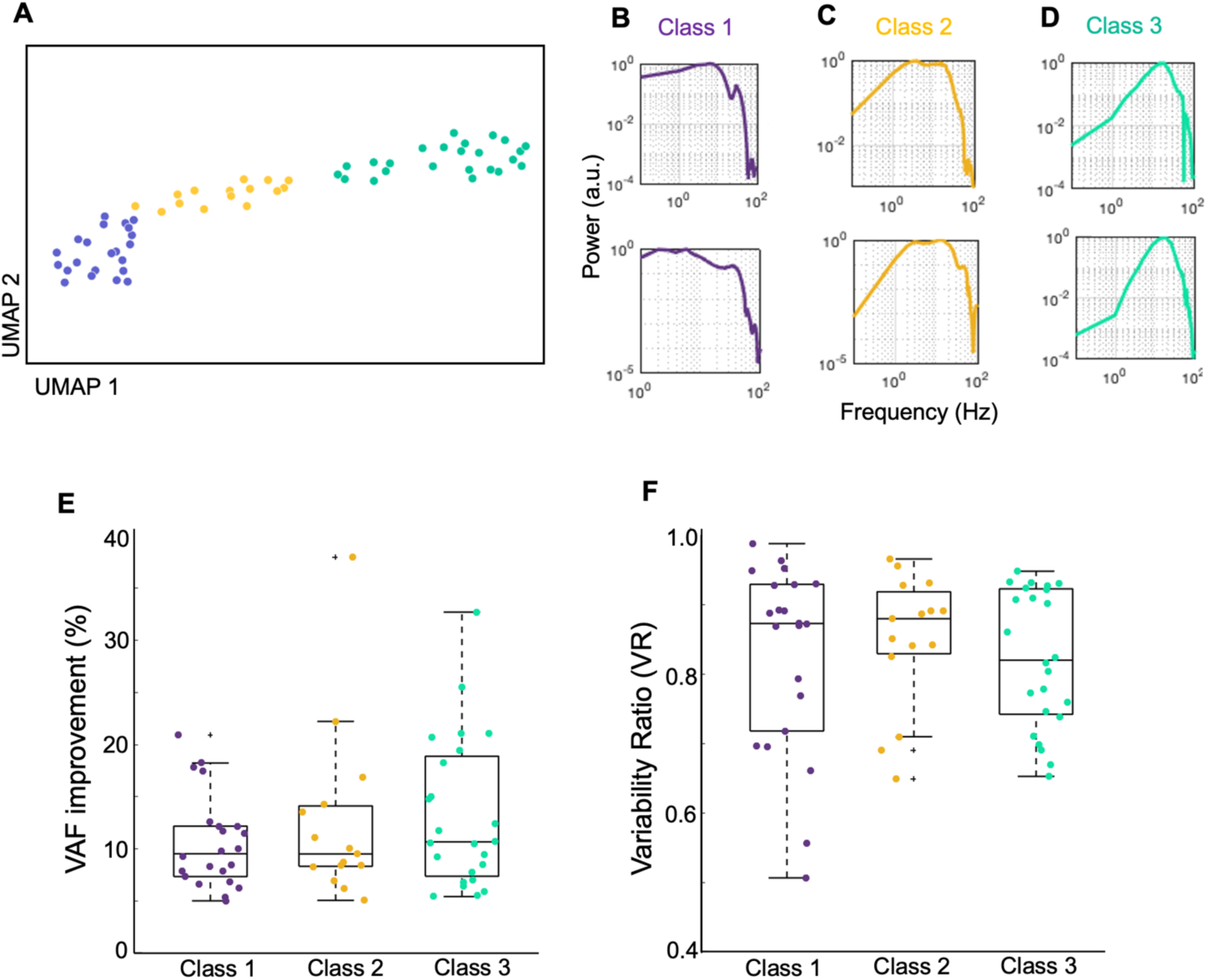
Diversity of filtered LFP power spectra. **A)** Scatter plot of filtered LFP power spectra in UMAP space, colored by Louvain cluster membership. Neurons with a VAF improvement > 5% (n=61) were used in this analysis. **B)** Two example power spectra for class 1 (purple cluster in panel A). **C)** Similar to B, for class 2 (yellow cluster in panel A). **D)** Similar to B and C, for class 3 (cyan cluster in panel A). **E)** VAF improvements of neurons which belong to the 3 classes. **F)** Similar to C, for the variability ratios of neurons.

In the above analysis, only the LFP from one nearby channel was used in the model. We evaluated further improvements in VAF by including LFPs from all the channels in grey matter (except the primary channel for the recorded neuron). Re-training of this model including the cortical state-driven pathway with all LFP channels provided separate temporal filters for each channel. Estimated filter weights for the first example neuron (Fig 7E) had lower magnitudes (± 0.008) for the first neuron compared to the second neuron (Fig 7F, ± 0.03). The filter weights showed clear spatial distribution for the second neuron, i.e., having higher magnitude weights at recording sites distributed around the primary channel for the neuron (red dashed line in Fig 7F). This spatial distribution of estimated weights was not observed for the first neuron (Fig 7E). The model prediction accuracy (VAF) improved by 2-fold for the second neuron (VAF 5.23% to 11.55%) while negligible improvement was observed for the first neuron (VAF 48.85% to 49.90%). Thus, these results support the idea that the variable portion of a neuron’s response might be caused by cortical state dynamics which are reflected in the LFP. Further, the second neuron which had a higher *VR* benefited more from the LFP-driven pathway, while showing strong coupling of the spiking response to ongoing network activity across nearby channels.

#### Contribution of MUA-based cortical state to model performance

Previous work has shown that average firing rate of a population of simultaneously recorded nearby neurons can be predictive of the observed trial-to-trial response variability of individual neurons (Okun et al., 2015). We evaluated whether the predictive performance for a single neuron’s spiking activity improves by incorporating MUA-based cortical state dynamics measured from two nearby channels (i.e., 100 μm above and below) in an additional parallel pathway in our model. MUA from the same channel was not used, to avoid potential contamination from the recorded neuron itself. Further signal “cleaning” was carried out by removing co-occurring spike signals in adjacent channels to avoid the effect of spikes from the same neuron appearing in multiple nearby channels (see Methods; S3 Fig).

Alternatively, MUA signals from the same channel could have been used after removing the spikes from the target neuron. However, to keep the processing steps comparable to LFP, we removed MUA signals from the same channel. Preliminary analysis showed the sufficiency of using only two MUA channels - the predictive accuracy with just two channels was higher (or similar) compared to that using additional channels. The MUA filter weights located further from two channels were close to zero in magnitude (S3 Fig), so including them in the model can cause the predictive accuracy to drop due to the increase of the number model parameters. Therefore, we used MUA signals from only two nearby channels separated by 100 μm from the neuron’s recorded channel and re-trained the model with the stimulus-driven pathway along with the pathway containing LFP signals from all the channels.

The pair of MUA temporal filters from two channels for the first example neuron (Fig 7G) were very small (+0.006 / -0.004) compared to those for the second neuron (Fig 7H) (+0.04/ -0.02). The temporal filters showed prominent peaks at latencies around 20-30 ms for the second neuron, but this effect was not observed for the first neuron. The VAF accuracy further improved for the second neuron from 11.55% to 17.41%, while negligible improvement was observed for the first neuron (VAF 49.90% to 49.45%). Thus, the second example neuron benefited comparatively more from inclusion of the MUA-driven pathway, similar to the effect from incorporating the LFP-driven pathway. Estimated LFP and MUA temporal filters for additional example neurons are shown in S9 Fig.

#### Improvement in estimated spatiotemporal receptive fields

Given the observed VAF improvement for the second example neuron compared to that for the first, we examined whether we could also observe parallel improvements in the estimated receptive fields. We evaluated the spatiotemporal receptive fields (i.e., restorations produced by convolving the spatiotemporal filter weights by the Gaussian map), while progressively incorporating the cortical state effects from both LFP and MUA signals.

Fig 9 shows, for the two example neurons, estimated receptive fields when using models with the stimulus-driven pathway alone (base model) in the first row, with LFPs from a nearby channel (Model 1) in the second row, with LFP from all the channels (Model 2) in the third row, and with the full model incorporating LFP from all the channels and MUA from two nearby channels in the fourth row (Model 3). For the first example neuron, we did not observe qualitative improvements in the receptive field (Fig 9A), which was not surprising in view of the absence of significant improvements in predictive performance (VAF=48.85% with the stimulus-driven pathway alone and only VAF=49.45% with the full model). However, for the second example neuron, we observed clear qualitative improvements in the receptive field after incorporating the cortical state effects (Fig 9B). This is evident from both the clear localization of the inhibitory and excitatory regions in the peak time lag at -26.7 ms, and better delineation of excitatory and inhibitory regions at the larger time lags (e.g. appearance of temporal reversal of the receptive field at time lag of -66.7 ms), after incorporating LFP- and MUA-driven pathways rather than simply using the stimulus-driven pathway (Fig 9B). This is also evident from the z-score values for the receptive fields of the two example neurons at the peak time lag. Neuron 1 had a z-score of 1.13 from the base model and 1.17 from model 3 (z-score improvement=0.04), while neuron 2 had a z-score of 1.55 from the base model and 1.79 from model 3 (z-score improvement=0.24). These examples suggest that for neurons with higher variability, accounting for the cortical state dynamics can not only improve the model predictive accuracy (VAF), but provide better visualizations of the spatial receptive fields and their temporal dynamics.

**Fig 9:**
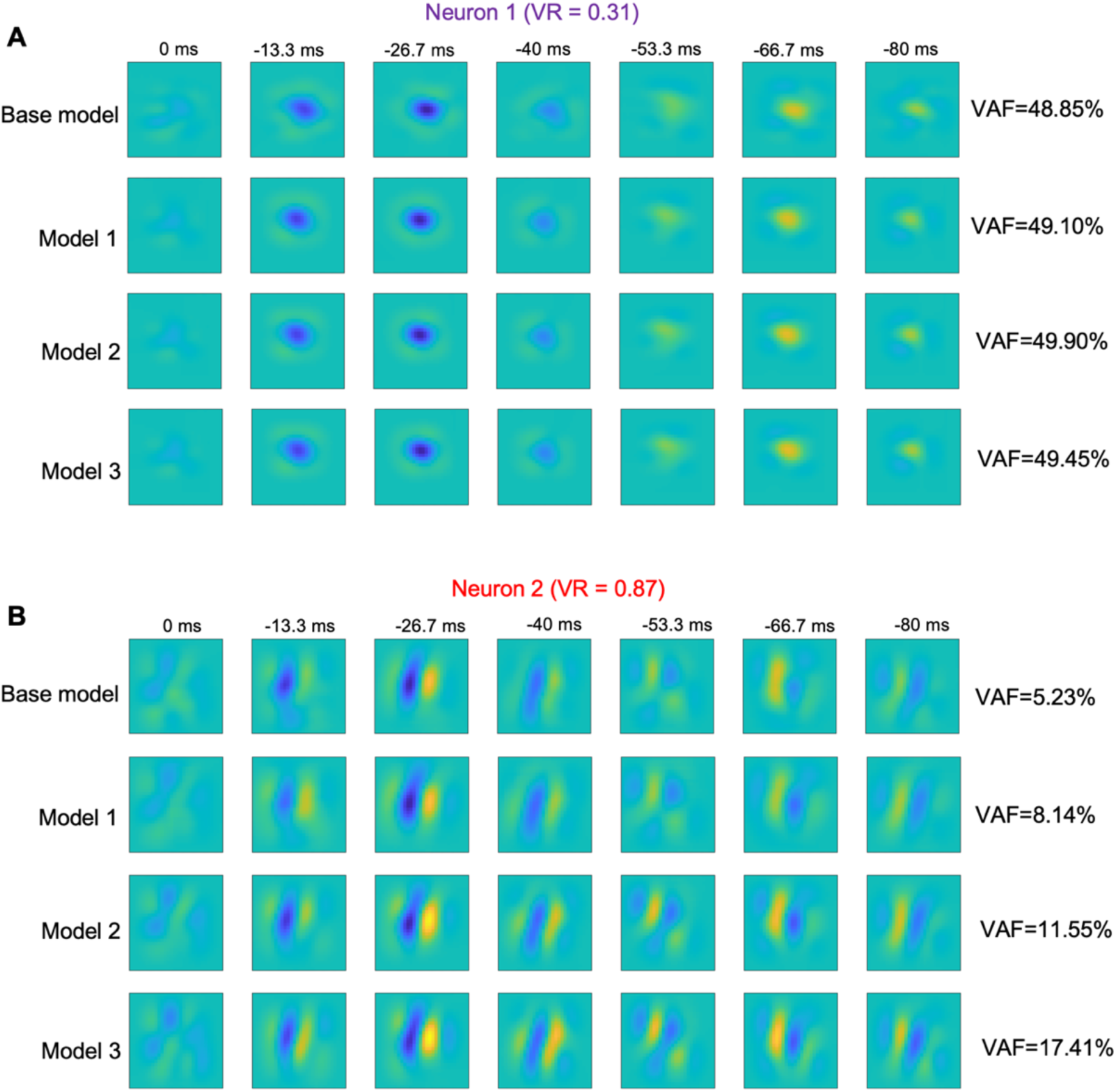
Improvements in receptive field estimates with cortical state. **A)** Improvement of spatiotemporal receptive fields with the incorporation of the brain state effects for example neuron 1. The base model is the system identification model with only the stimulus-driven pathway, Model 1 incorporates cortical state effects from one LFP channel located 100 μm away, Model 2 incorporates all the LFP channels and Model 3 additionally incorporates two nearby MUA channels. **B)** Similar to A, for example neuron 2.

#### VAF and receptive field improvements across the population of neurons

The initial VAF of the n=154 neurons with only the stimulus-driven pathway (base model) ranged from 0% to nearly 50%. The improvements in model predictive power for this sample population of neurons is shown in Fig 10. With the stimulus-driven pathway alone (base model), neurons exhibiting a lower variability ratio (*VR*) yielded higher predictive performance, whereas those with a higher variability exhibited lower predictive performance (Fig 10A). This inverse relationship between VR and the VAF predictive performance was confirmed statistically (Fig 10A, r=-0.89, p=2.29e-54).

**Fig 10:**
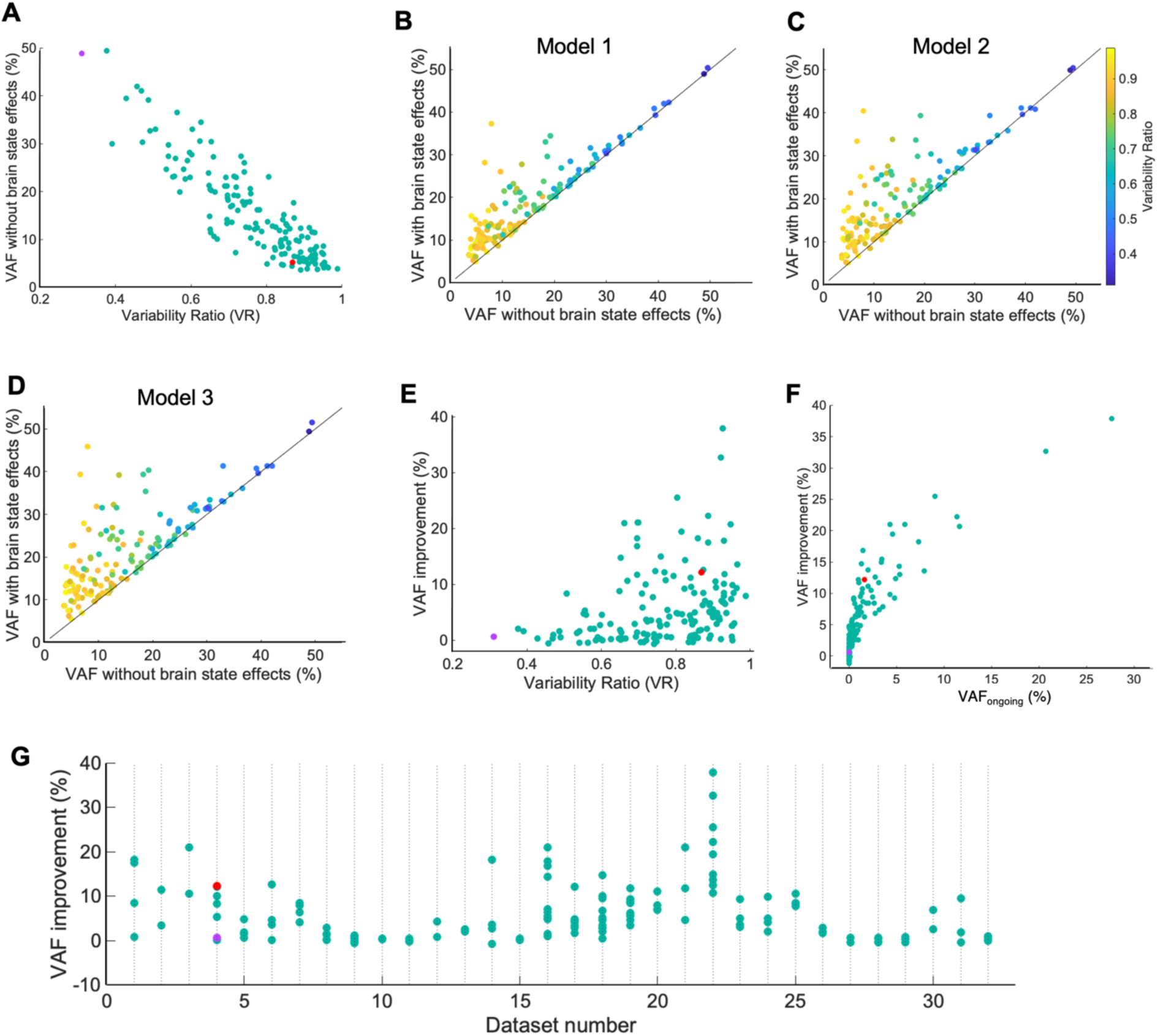
Improvements in predictive performance for the models with cortical state indicators, across the population of recorded neurons. **A)** Scatter plot for the population of neurons (n=154) showing the VAF from the stimulus-driven pathway against response variability ratio (*VR*). **B)** Scatter plot for the population of neurons showing VAF from model with LFP from the above channel, against VAF from only the stimulus-driven pathway. Line indicates 1:1 ratio. Data points for each neuron are color coded according to their variability ratio. **C)** Similar to B, with VAF from the model with LFPs from all channels. **D)** Similar to B, but showing VAF from model with LFPs from all channels and MUA from above and below channels. **E)** Improvement in VAF after incorporating cortical state effects in Model 3, plotted against response variability ratio for each neuron. **F)** Improvement in predictive VAF plotted against the *VAF_ongoing_* for each neuron. **G)** Distribution of VAF improvements of neurons across 32 datasets, each containing at least 2 simultaneously recorded neurons. Each data point on a column shows a simultaneously recorded neuron in that dataset.

Given the very different observed VAF improvements in two example neurons (Fig 9), we examined the improvements of VAF across the population of 154 neurons. We analyzed the gradual improvements in the VAF predictive accuracy when we incorporate LFP signals from a channel located 100μm away (Model 1, Fig 10B), LFPs from all the channels (Model 2, Fig 10C), LFPs from all the channels and also MUA signals from two nearby channels (Model 3, Fig 10D), compared to the base model (stimulus-driven pathway alone, abscissa of plots in Fig 10B,C,D). The data points of these scatter plots represent single neurons, color-coded according to variability ratio. These neurons showed statistically significantly higher VAFs compared to the base model using LFPs from a nearby channel (Fig 10B; VAF from Model 1 compared to base model: p=9.53e-05, n=154), LFPs from all the channels (Fig 10C; VAF from Model 2 compared to base model: p=8.34e-07, n=154) and with the complete model including both LFP and MUA parallel pathways (Fig 10D; VAF from Model 3 compared to base model: p=9.95e-09, n=154). In addition, we observed successive improvements in VAF from model 1 to model 3 (median VAFs for base model= 12.44%; model 1= 17.17%; model 2= 18.67%; model 3= 19.82%; n=154).

These improvements were greater for neurons with higher response variability, but relatively minor for neurons with a lower response variability (Fig 10E Pearson’s r=0.49, p=1.52e-10). Overall, we observed improvements of the predictive performance (VAF) in the population of neurons improving from 0% to 40% (Fig 10E). Given the range of observed VAF improvements, we explored whether they are related to the amount of variability explained by our model. For this, we calculated the measure “ *VAF_ongoing_*” in which we compare, for each neuron, the variable part of the response to the additional response predicted by Model 3 compared to the base model (see Methods). Higher *VAF_ongoing_* indicates that the additional responses predicted by Model 3 compared to the base model are accounting for most of the variable part of the neuronal response. The observed overall VAF improvements correlated well with the *VAF_ongoing_* values, i.e., the neuronal response variability was predicted well by our cortical state-driven pathways (Fig 10F: r=0.85, p= 1.65e-44). Thus, these results illustrate the capability of the model to account for the observed trial-to-trial response variability by incorporating cortical state dynamics along with the stimulus-driven pathway.

Simultaneously recorded neurons in the population often exhibited quite different amounts of response variability (Fig 4E). Therefore, we explored whether these simultaneously recorded neurons also show different amounts of VAF improvements after accounting for the cortical state effects using LFP and MUA signals. Across the sets of simultaneously recorded neurons, we observed different amounts of VAF improvements ranging from a maximum difference of 27.2% to a minimum of 0.07% with a median of 4.39% (Fig 10G).

Given the observed apparent improvements in the receptive fields of the second example neuron (Fig 9B), we examined the extent of this improvement across our population of neurons. To quantify the apparent qualitative improvements in estimated receptive fields for the population of neurons, we calculated z-score values for the receptive fields at the peak time lag (i.e., -26.7ms; see Methods). The z-score values after incorporating cortical state dynamics (median 1.08, inter-quartile range 0.86) were slightly larger than those from the base model with the stimulus pathway alone (median 0.91, inter-quartile range 0.72), and the difference was statistically significant (Fig 11A, p=1.38e-02, n=154). Fig 11B plots the z-scores for the two models plotted against one another. Among the 154 neurons, 100 neurons (64.94%) have higher z-scores for the full model (dots above the diagonal line), while 54 neurons (35.06%) have lower z-scores for the full model (dots below the diagonal line). However, Fig 11C shows that the improvements in the receptive fields were not statistically correlated with the observed VAF improvements (r=0.13, p=9.6e-02).

**Fig 11:**
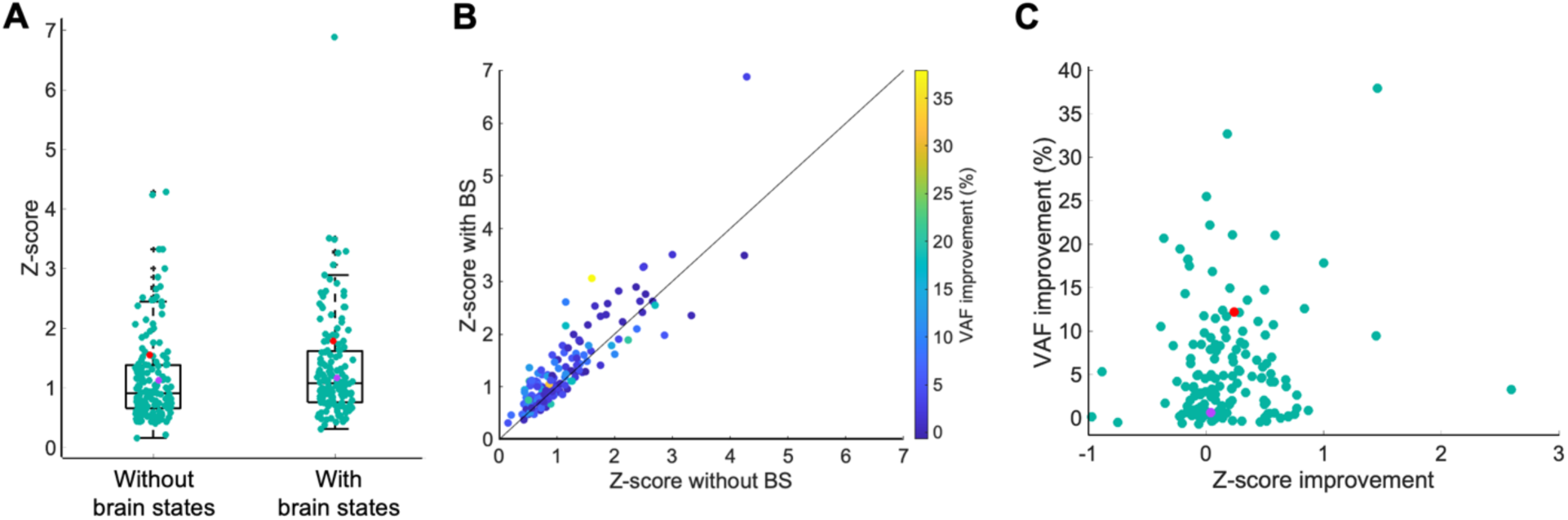
Receptive field improvements with cortical state across the sample population of neurons. **A)** Box plot of z-scores for the receptive fields estimated without and with cortical state-driven pathways. **B)** Similar to A, shown as a scatter plot. **C)** Scatter plot of z-score improvement and VAF improvement.

## Discussion

A major percentage of responses recorded from primary visual cortex neurons cannot be explained as due to visual stimuli alone, due to trial-to-trial response variability. The present study evaluated the hypothesis that the observed response variability arises from cortical state fluctuations. To test this idea, we proposed a compact convolutional neural network model of each neuron’s visual receptive field, that also incorporated temporally filtered LFP and MUA signals as measures of cortical state dynamics (Fig. 3). The fitted model showed significantly improved accuracies in predicting neuronal responses and in estimating receptive fields, compared to a model with a stimulus-driven pathway alone (Fig 10B-D, Fig 11). The improvements in predictive performance were greater for neurons with higher trial-to-trial response variability than for those showing more reliable responses across trials (Fig 10E-F). These results support the idea that LFP and MUA signals carry information about response variability caused by the fluctuations in cortical states (Cui et al., 2016; Kelly et al., 2010).

Furthermore, neurons recorded simultaneously could evidently be affected differentially by cortical state variations. Some of the simultaneously recorded neurons showed substantially higher response variability across trials and higher predictive accuracy improvements with incorporated cortical state effects in the model, while other neurons produced relatively reliable responses across trials and showed lower predictive accuracy improvements (Fig. 4E, 10G). In addition, the estimated temporal filters for LFP and MUA were neuron-specific, and filter different frequencies of the LFP signal, demonstrating an important factor in how different neurons are affected differentially by cortical state dynamics (Fig. 7, S6, S9 Figs).

### Inference of cortical state fluctuations

Our evaluation of the effects of cortical states on response variability is based on the prevalence of ongoing cortical state dynamics within a data recording session. We confirm this by deriving a global fluctuation index (GFI, adapted from (Schölvinck et al., 2015)) from binned MUA during the blank inter-sweep periods, and a synchrony index (SI, adapted from (Spacek & Swindale, 2016) from LFPs across each data recording session. These fluctuations showed a similar pattern when compared at the same time points (Fig 4, red dots in panel E vs. panel C data points, S8 Fig). Thus, even though the animals are under anesthesia, their cortical state fluctuated continuously, consistent with the idea of states varying in a continuum (Harris & Thiele, 2011).

The estimates of cortical state could constrain the performance of our proposed model. We used simultaneously recorded MUA and LFP brain signals to quantify cortical state dynamics. Since MUA and LFP signals contain information in relatively high and low frequency ranges, respectively, we expected that they might carry complementary or independent information about cortical states. Cortical state characteristics have often been regarded as slow fluctuations on a time scale of seconds or more (Harris & Thiele, 2011) - these characteristics should be reflected in low frequency LFP signals. However, our estimated temporal filters for both LFP and MUA contain an optimal time window in the millisecond scale (Fig. 7). Preliminary analysis showed that increasing the temporal window length by including more time points or including an additional parallel pathway for ultra-low LFP signals (<0.5Hz) did not bring additional improvements in predictive accuracies. Therefore, our millisecond scale time window may be primarily capturing rapid up-down phases (Harris & Thiele, 2011; Munro et al., 2015) embedded within the slow fluctuations. These relatively fast dynamics could also reflect millisecond-scale resolution of brain state embeddings, including substates and rapid events shown in recent studies (Parks et al., 2023). In spite of similar time scales of the estimated filters, incorporating the MUA pathway does yield improved predictive performance, suggesting that it provides some additional higher temporal resolution information related to cortical states.

### Variation in LFP frequencies of importance

Previous studies showed that different frequency bands of EEG and LFPs correspond to distinct behavioral states (Saby & Marshall, 2012; Yun et al., 2023). For example, low frequency delta signals are associated with unconsciousness or deep-dreamless sleep, and high frequency gamma signals are associated with motor functions and higher mental activity (Saby & Marshall, 2012). Across our sample of neurons, we observed a wide diversity in the estimated LFP filters, and consequently of the LFP frequencies effective for the cortical state-driven pathway (Fig 7A-D, S6, S9 Figs). This result suggests that cortical state dynamics can affect individual neurons differentially.

The filtered LFP frequency bands from the system identification model aligned well and showed substantial similarities to values of phase locking strength computed using LFP signals and spiking responses (i.e. similar frequency ranges in Fig 2B, C vs Fig 7C, D and in S9 Fig B vs C). This validated the capability of our model in estimating temporal filters to extract the significant LFP frequency bands when the whole LFP signal provided as an input to the model. Therefore, our approach works flexibly without requiring the use of LFP signals segmented into pre-determined frequency bands.

Previous studies in area MT of awake behaving macaque monkeys showed strong links of spiking activity to specific frequency bands of LFPs, including high frequency gamma (30-70 Hz) and low frequency delta (1-4 Hz) bands (Cui et al., 2016). This difference from our results, in which we saw a range of relevant frequencies, could be due to various factors, including the difference of species, brain areas and brain states (awake vs under anesthesia), and our approach of estimating filters applied to raw LFPs rather than wavelet coefficients.

### Cortical state-driven pathways

Our model architecture for cortical state pathways utilizes simple linear filters that are unique for individual neurons, and to infer cortical states from LFP and MUA signals. We evaluated several alternative approaches including filter-rectify-filter (FRF) cascades to capture envelopes of high frequency LFPs, using dimensionality-reduced LFP signals, and allowing the brain state modulation to act at intermediate stages along the visual processing pathway. However, the predictive accuracy of each of these was comparatively lower. Nevertheless it is conceivable that some more complex model architecture might have better incorporated these population signals, though at the expense of additional model parameters.

Another limitation is that we measured LFPs and MUA only for sites along our linear array, but signals from other columns and brain areas might carry relevant and distinct cortical state-related information. Including these additional signals, e.g. in parallel pathways in a model like ours, might further improve the prediction accuracies.

Other studies have used measures of cortical states derived from other brain signals including scalp EEG (Hobson & Pace-Schott, 2002) and hippocampal LFPs (Pedrosa et al., 2022). Cortical state can also be reflected in behavioural signals such as whisker movements (Poulet & Petersen, 2008), locomotion velocity and pupil diameter (McGinley et al., 2015). Including such signals in parallel cortical state-driven pathways could enrich a model’s representation of brain state fluctuations. The extent to which these more elaborate models could boost the neural response predictive accuracy is uncertain since many of these brain state-indicative signals may correlate with one another (Larsen & Waters, 2018). Nevertheless, it would be interesting to examine whether using these behavioural signals in additional parallel pathways in a model like ours could further improve the predictive accuracies.

### Spike contamination in LFP signals

A potentially important concern is that LFPs obtained from a electrode recording site could be contaminated by spikes from nearby neurons. Therefore, if we use the LFP from a given channel to predict the spiking activity of a neuron recorded from the same channel, that LFP signal could contain a significant contribution from the spike itself and yield misleading predictive accuracy.

While previous work showed that although spikes can be removed from LFP using a complex spike removal algorithm (Zanos et al., 2011), residual spike contamination effects may still remain (Pesaran et al., 2018). Hence one remedy is to use spikes and LFPs from nearby but distinct recording sites (Pesaran et al., 2018). Consistent with this, (Cui et al., 2016) found similar system identification model weights estimated using spike-removed LFPs versus LFPs 100 μm away from the spike recording electrode. Consequently, we excluded LFPs from the same channel as that of the recorded neuron, to mitigate effects of residual spike signals from that neuron.

### Issues in accounting for response variability

The present model did not completely account for the observed response variability (Fig. 10F). Even with the incorporation of brain state-related signals in our model, the predictive performance remains well below 100% (Fig 10D). An elementary reason for predictive error could in part due to intrinsic neuronal noise, including cellular, electrical, and synaptic noise that arise at different stages of sensory processing (Faisal et al., 2008). Most sources of intrinsic noise would be relatively uncorrelated across neurons, and thereby contribute less to neuronal population signals. Hence, our model might not capture intrinsic noise as compared to brain state fluctuations that can exert significant effects on neuronal spiking variability. In addition, variability can arise due to stimulus-dependent factors including stimulus onset (Churchland et al., 2010), stimulus features including orientation and contrast or (Kohn & Smith, 2005), adaptation from previous visual stimuli (Snow et al., 2017). More elaborate model architectures, and/or additional brain state indicators, might be required to account for these effects.

Previous studies have found that neurons’ response variability increases across the visual hierarchy from retina to LGN to V1 and other higher cortical areas, and is also inversely correlated with the average firing rates across these three brain regions (Goris et al., 2014; Kara et al., 2000). Consistent with this, we did observe a negative correlation between the firing rates and response variability of neurons, which was relatively weak though statistically significant (Fig 4C). The observed weak correlation arise because our population contains only area 17/18 neurons, but not recordings across the visual hierarchy. However, if this correlation were greater, addition of a firing rate dependency into the model architecture might be useful.

### Stimulus-driven pathway

The stages in our stimulus-driven pathway are easily interpretable and applicable to both simple and complex cells in the primary visual cortex (Nguyen et al., 2024). However, important improvements in predictive performance might be gained by developing more elaborate model architectures to handle other known aspects of visual processing, particularly nonlinearities such as gain control (Ohzawa et al., 1985), surround suppression, adaptation, or second-order processing (Li et al., 2014). Some studies have reported that deep neural networks, e.g. with more filters and/or filter layers, can provide improved predictive power for cortical neurons’ visual responses (Cadena et al., 2019; Ukita et al., 2019; Zhang et al., 2019). While such approaches might be desirable and interesting, they would entail greater challenges in optimizing the additional model parameters while avoiding overfitting, and in the case of deep neural networks, would entail a substantial cost in interpretability.

Previous studies reported that the structures of V1 receptive fields could substantially change when measured in different contexts, e.g. with receptive fields becoming wider in synchronized cortical states and narrower during desynchronized states (Wörgötter et al., 1998). Preliminary analysis with a subset of our neurons revealed that in some cases the receptive field size increased with synchronized states, consistent with the (Wörgötter et al., 1998) report, while others exhibited an opposite result - in any case, these effects were relatively small (S10 Fig). Therefore, in our system identification approach we treated the receptive fields as having a common structure and size throughout a given data acquisition session, regardless of cortical state variations. If such changes were shown to be a more serious effect, then this limitation could conceivably be addressed in future approaches by using a model architecture that incorportates cortical state effects inside the stimulus-driven pathway.

### Improvements in receptive field estimation

Incorporating brain state fluctuations often led to improvements in predictions of neurons’ responses, and also to better estimates of RFs in terms of z-scores – however, across the sample population these two kinds of improvement were not statistically correlated (Fig. 11). This discrepancy could be due to multiple factors. The training of our model is based on optimizing the accuracy in predicting neural responses - in doing so, the optimization process might focus less on fitting the most accurate RF. Conversely, inadequacies of our visual pathway model, which would degrade the overall predictive accuracy, could vary substantially across neurons, in a manner not necessarily related to the z-scores. Thus for example, a poor visual response model applied to a neuron with little brain state dependence, might give a “clean” (high z-score) RF estimate which is quite biased, resulting in poor predictive performance.

In addition, our z-score measure might not be an ideal measure of the quality of an estimated RF. For example, collapsing the spatial z-score values into a single number (Eq 15) in deriving our z-score measure would not be completely appropriate for some neurons. As a result, comparing quantitative improvements in VAF to qualitative improvements in RFs in terms of z-score might lead to the observed discrepancies.

### Conclusions

Our results have demonstrated varying amounts of response variability, and of improvement in predictive performance, for different neurons - most interestingly, even for simultaneously recorded neurons. This heterogeneity of brain state effects across neurons places an important caveat on interpretations of global or large-scale (population) measures of brain state, such as scalp-recorded EEG. It would be interesting to further explore this heterogeneity, by examining whether there are systematic differences between neurons affected differentially by brain state, e.g. receptive field properties or laminar location.

## Acknowledgements

This work was supported by the Canadian Institutes of Health Research Grant MOP-119498 to C.L.B, and a Quebec FRQS doctoral fellowship to J.S. We thank Guangxing Li for technical support in experiments; and Amol Gharat and Phillipe Nguyen for contributions to data collection.

## Supporting Information

**S1 Fig.**
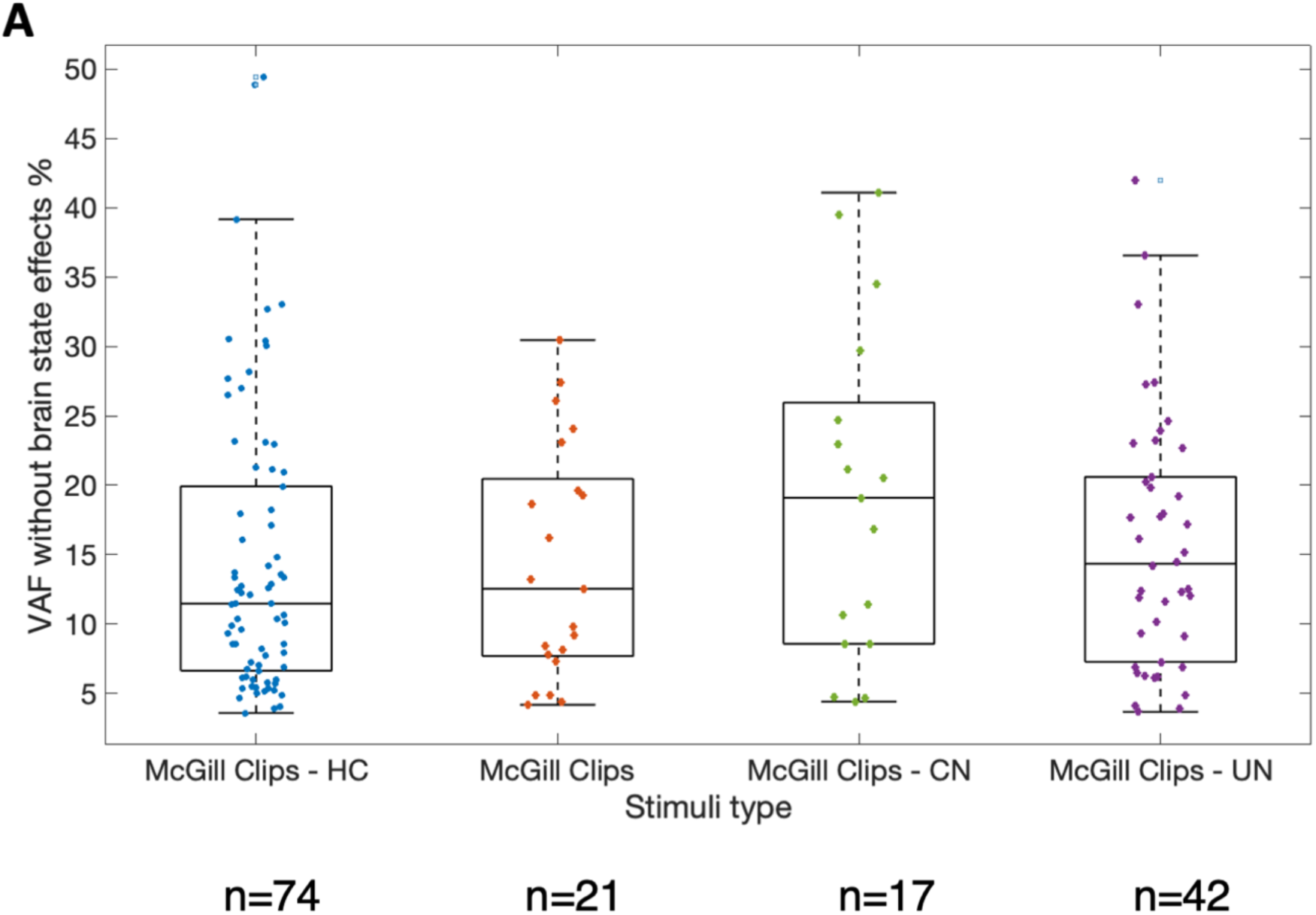
Comparison of prediction accuracies using different versions of natural image stimuli. Each data point represents a neuron’s VAF accuracy obtained from system identification model with the stimulus-driven pathway only for the four types of stimuli.

**S2 Fig.**
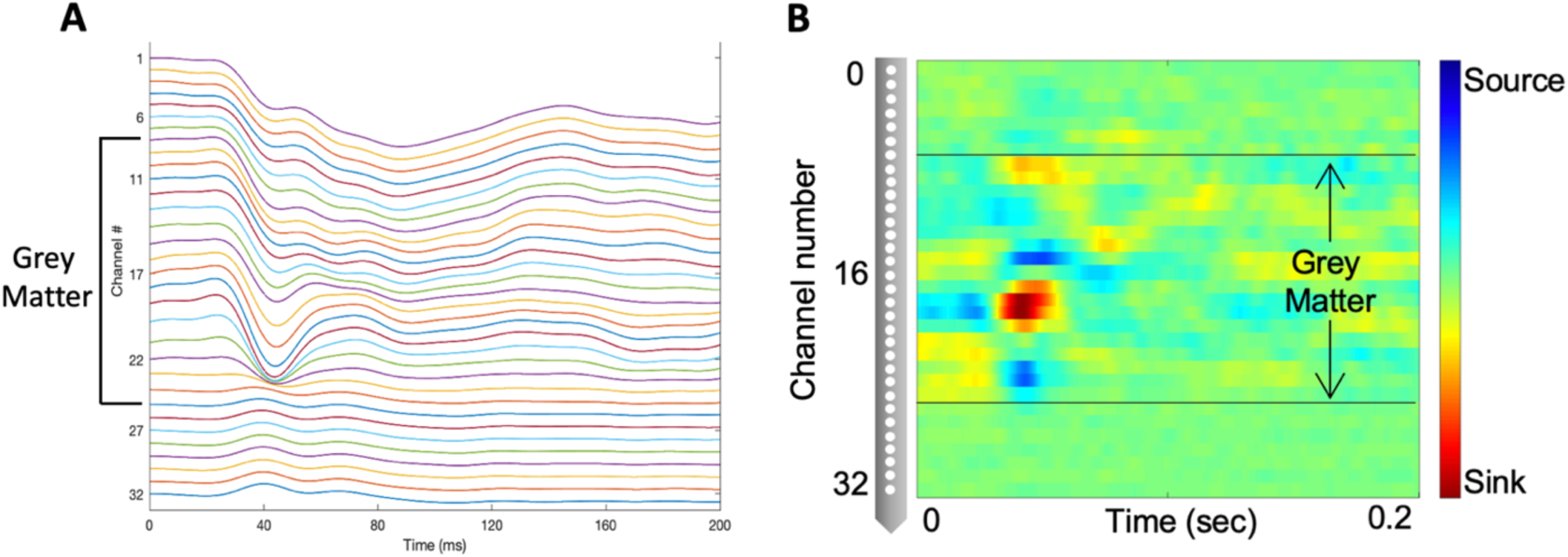
Identification of recording channels in grey matter. **A)** Raw LFP traces used to measure current source density (CSD) as a function of depth. We extracted LFP data (<100 Hz) from 32 channels while presenting sinewave gratings. We averaged the filtered LFP signals across trials to obtain the evoked response potential and used these to compute CSD. **B)** CSD plot generated using *CSD plotter* software (Pettersen et al., 2006). Recording channels were 100 μm apart, so the apparent gray matter extended across about 1800 μm. Channels with a prominent current sink in the CSD profile were specified as the granular layer, and sources above and below this sink were identified as the supragranular and infragranular layers respectively.

**S3 Fig.**
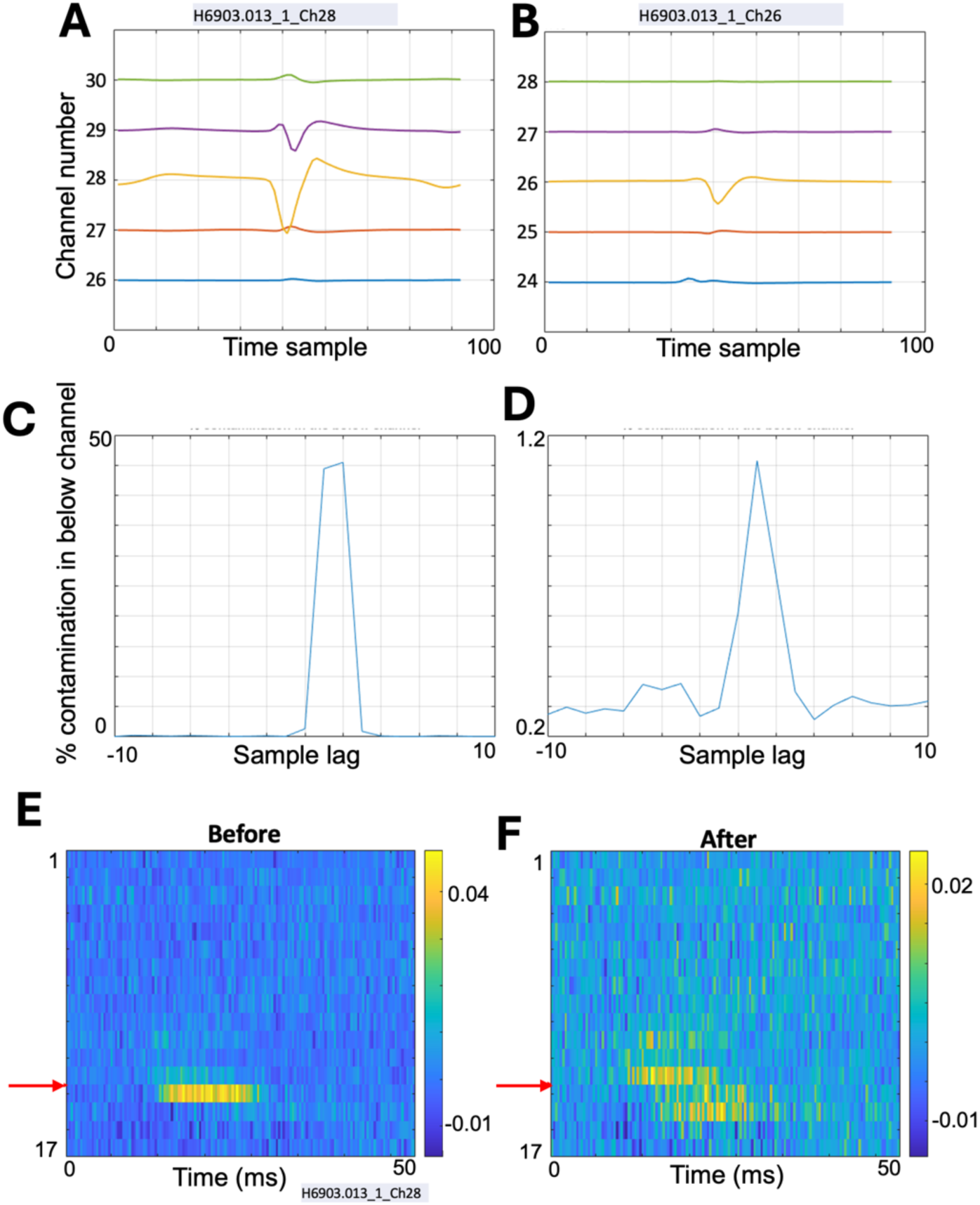
MUA cleaning. **A, B)** Spike waveforms recorded in adjacent channels for two example neurons. **C and D)** Percentage of spikes in the below channel contaminated with spikes of the neurons in A and B. To compute this, we count the number of spikes in the neighboring channel (i.e., channel located 100μm below, in this example) co-occurring with the neuron’s spikes from the primary channel (here, 28) from which it was recorded. If this count is >10% of the total spikes in the neighbouring channel, we remove those spikes from the neighbouring channel. **E)** Estimated filter weights from MUA pathway when MUA from all the channels is used as inputs before removing spike contamination from the neighbouring channels. **F)** Same as E after removing spike contamination from the neighbouring channels.

**S4 Fig.**
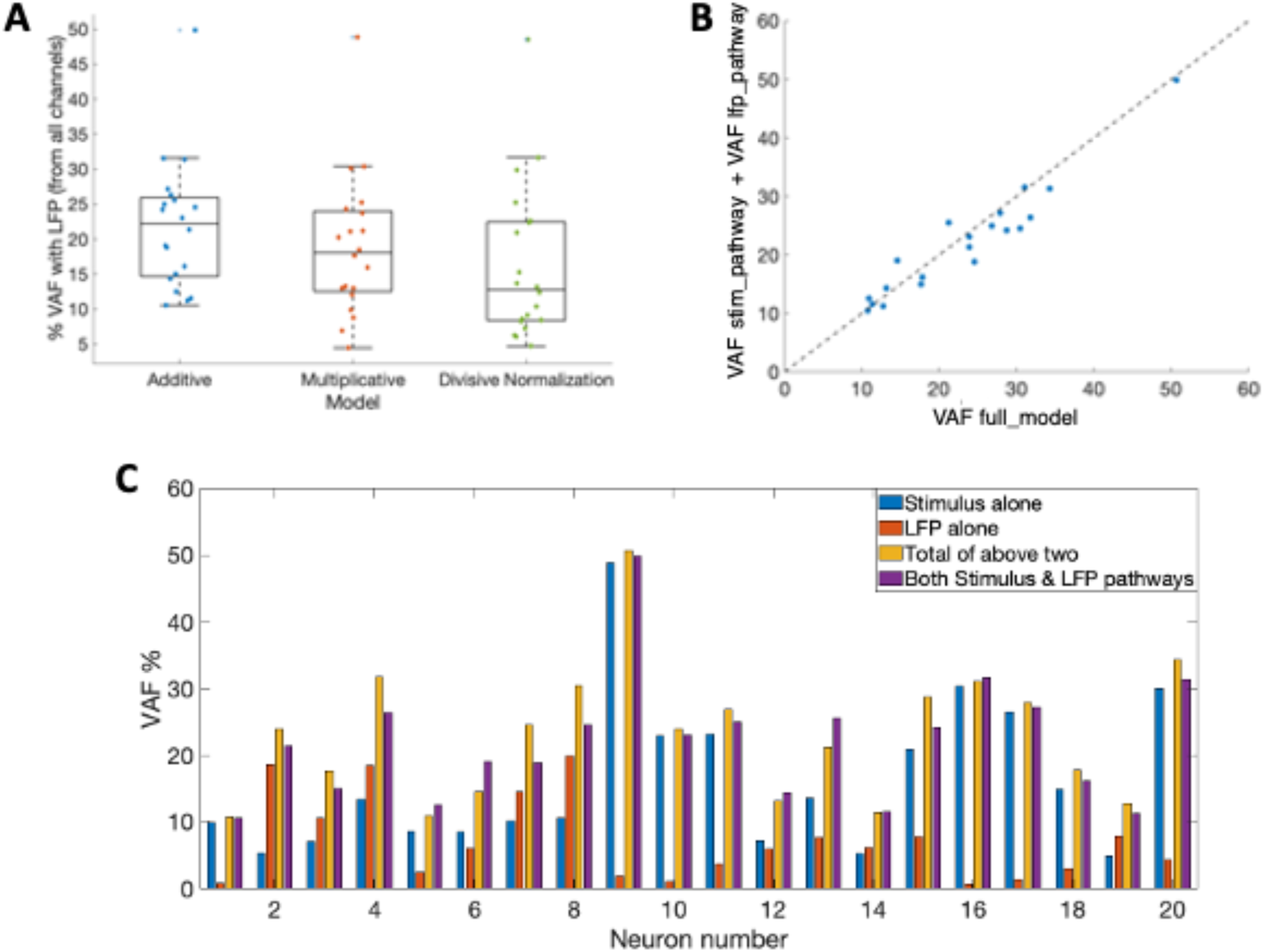
VAF additivity. **A)** Box plot showing VAF from model with LFP from all channels acting additively, multiplicatively, or with divisive normalization. **B)** Scatter plot showing the summation of VAFs from stimulus pathway alone (VAF stim_pathway) and LFP pathway alone (VAF lfp_pathway) vs VAF from full model containing both stimulus and LFP pathways (VAF full_model). Dashed line indicates 1:1 relationship. **C)** Bar graph showing VAFs from stimulus pathway alone, LFP pathway alone, summation of both, and VAF from the full model containing both stimulus and LFP pathways.

**S5 Fig.**
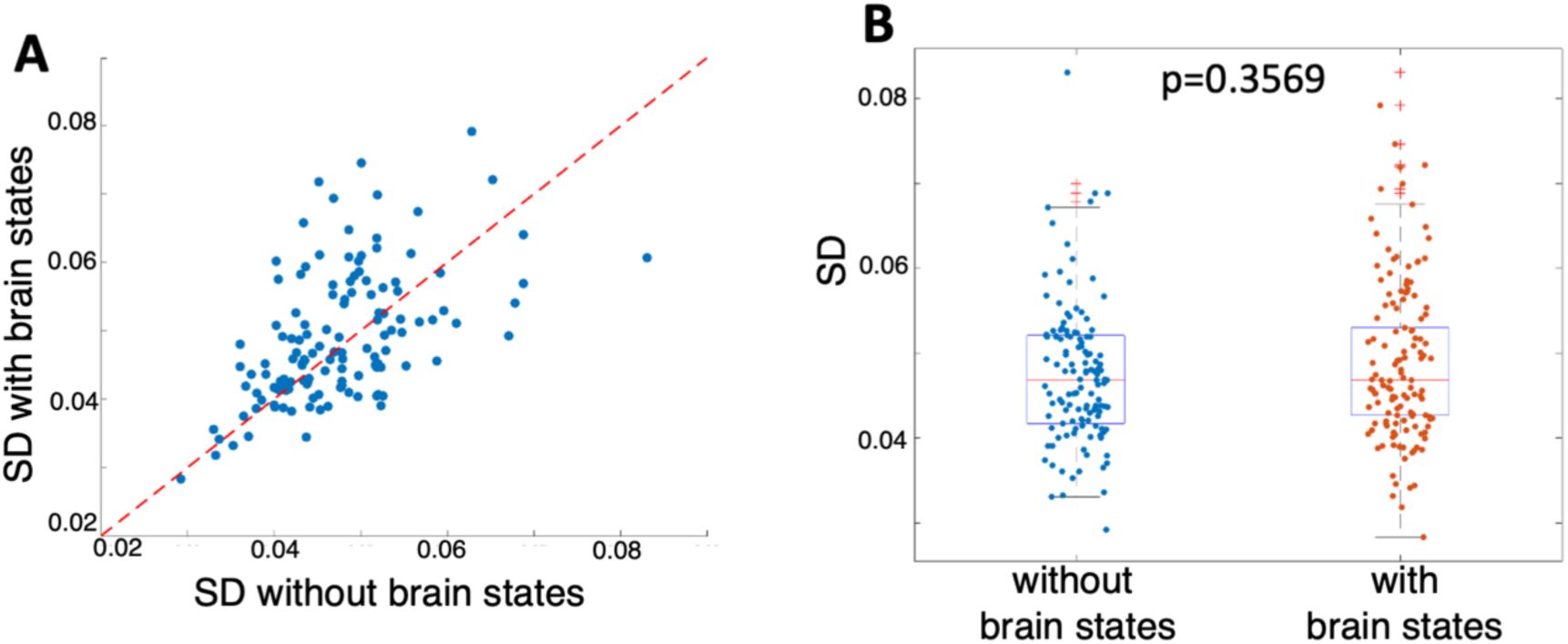
Z score calculation. **A)** Scatter plot showing standard deviation of the zero^th^ time lag of the estimated receptive field with brain state pathways, vs. the same but for the base model without brain state. **B)** Distribution of standard deviations of the zero^th^ time lag of the estimated receptive fields without, and with, brain state models.

**S6 Fig.**
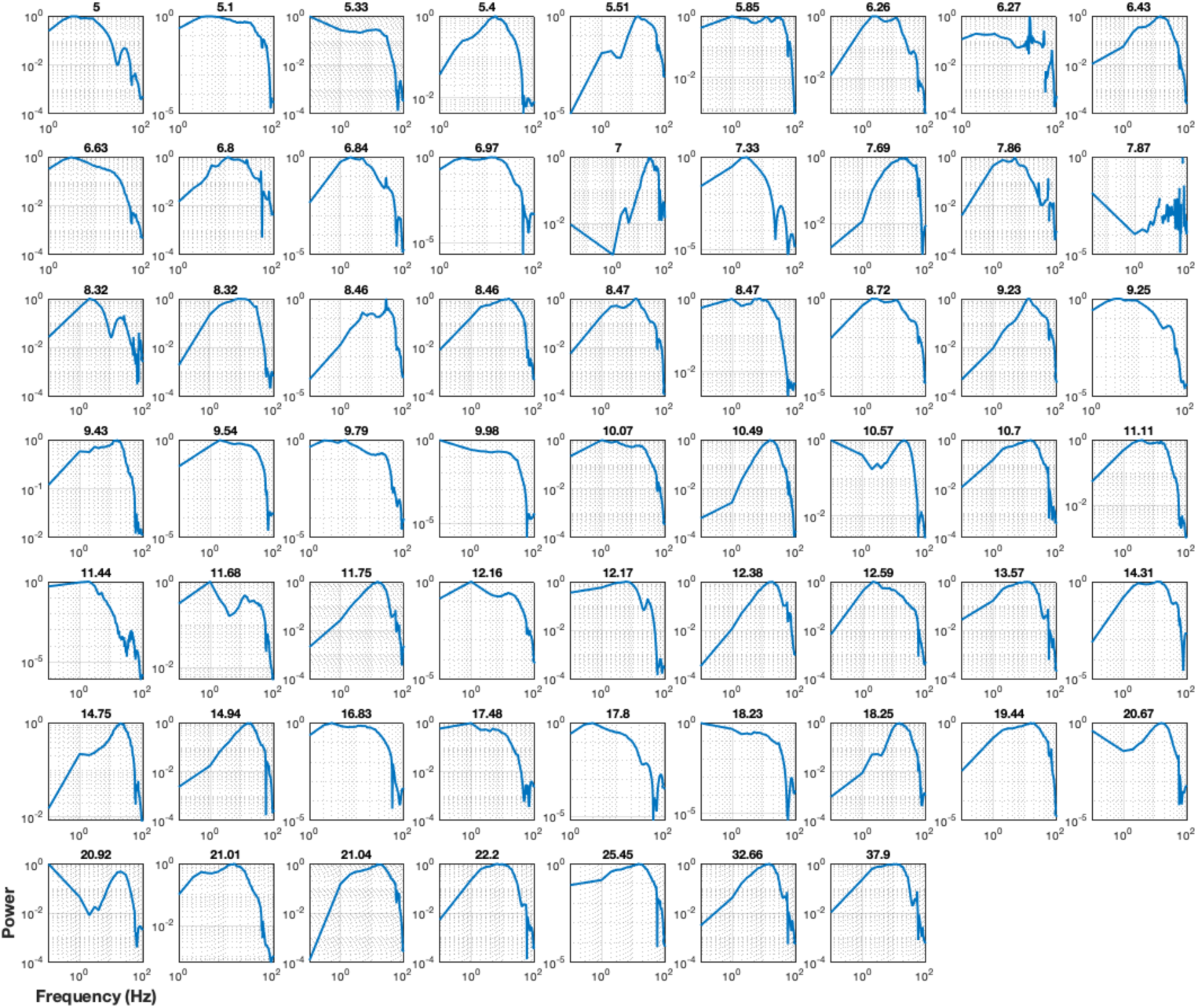
LFP frequencies of importance for different neurons. Power spectra of recorded LFPs filtered with estimated LFP temporal filters for neurons with >5% VAF improvements (n=61). VAF improvements are reported on top of each power spectrum.

**S7 Fig.**
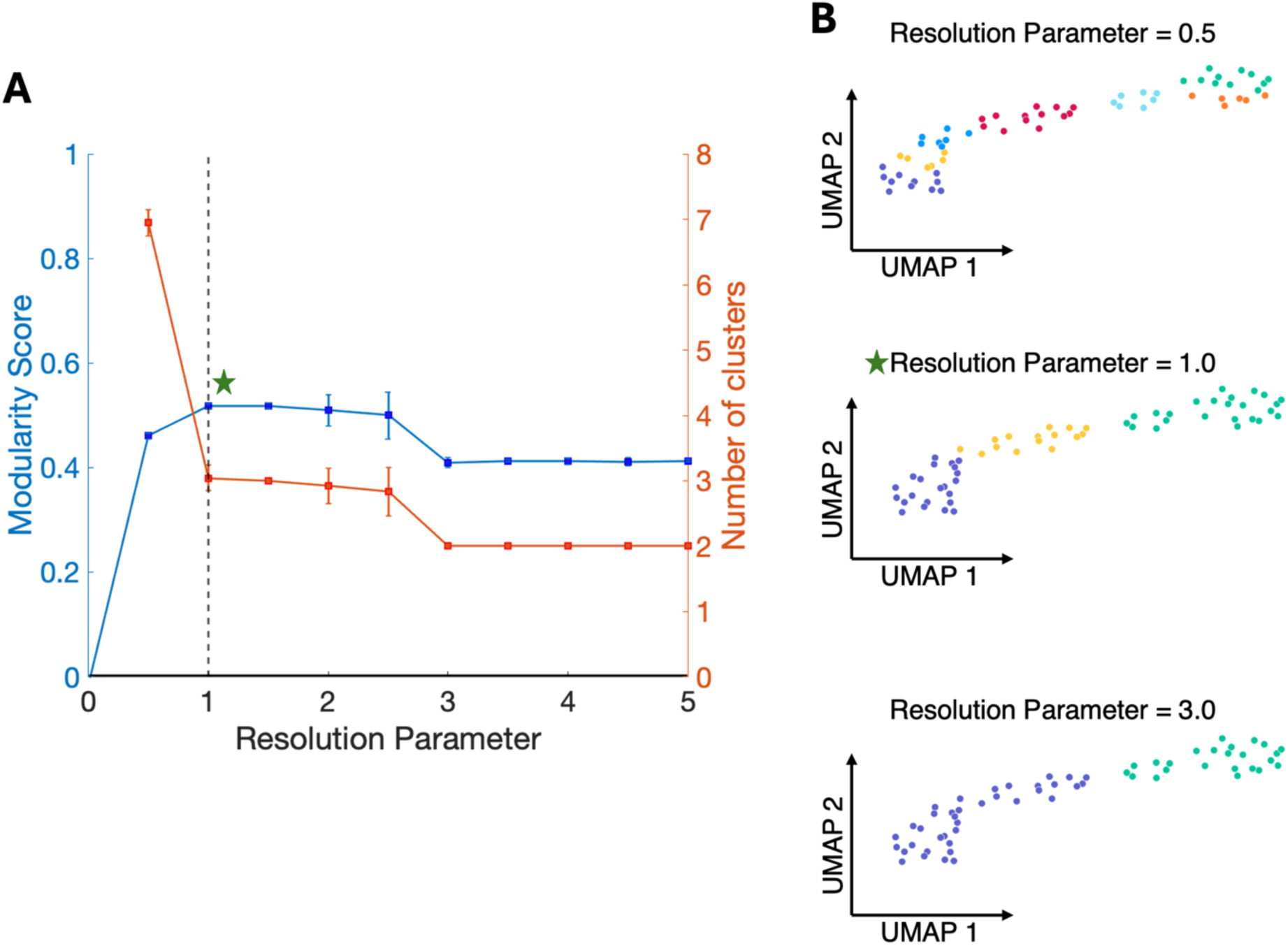
Clustering resolution parameter vs modularity score. **A)** Modularity score and number of clusters vs resolution parameter in Louvain clustering. The dotted line when the modularity score is the highest at resolution parameter = 1.0 was selected for clustering. **B)** Clusters obtained with resolution parameter = 0.5, 1.0 and 3.0. Green star shows the clustering at the selected resolution parameter of 1.0.

**S8 Fig.**
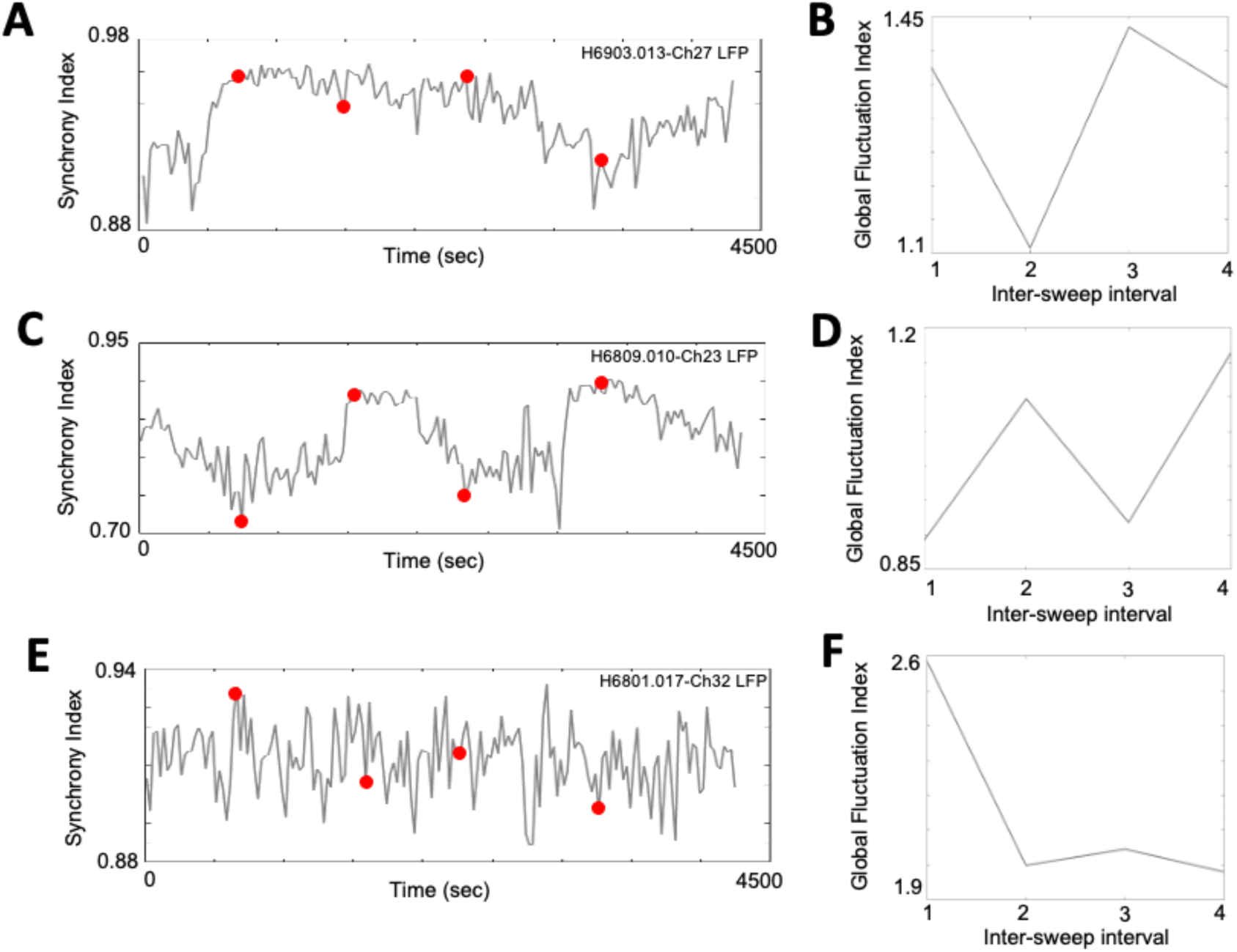
Cortical state fluctuations within data recordings. **A)** Synchrony index (SI) defined by equation 4 using PSD of a deep layer LFP signal. Red points indicate inter-sweep intervals. **B)** Global fluctuation index (GFI) defined by equation 3. **C, E)** Similar to A for another two datasets. **D, F)** Similar to B for another two datasets. The Pearson’s correlation between SI and GFI and its statistical significance for the three data recordings are r=0.65, p=0.35 (top row); r=0.99, p=5.9e-03 (middle row); and recording 3: r=0.98, p=1.7e-02 (bottom row).

**S9 Fig.**
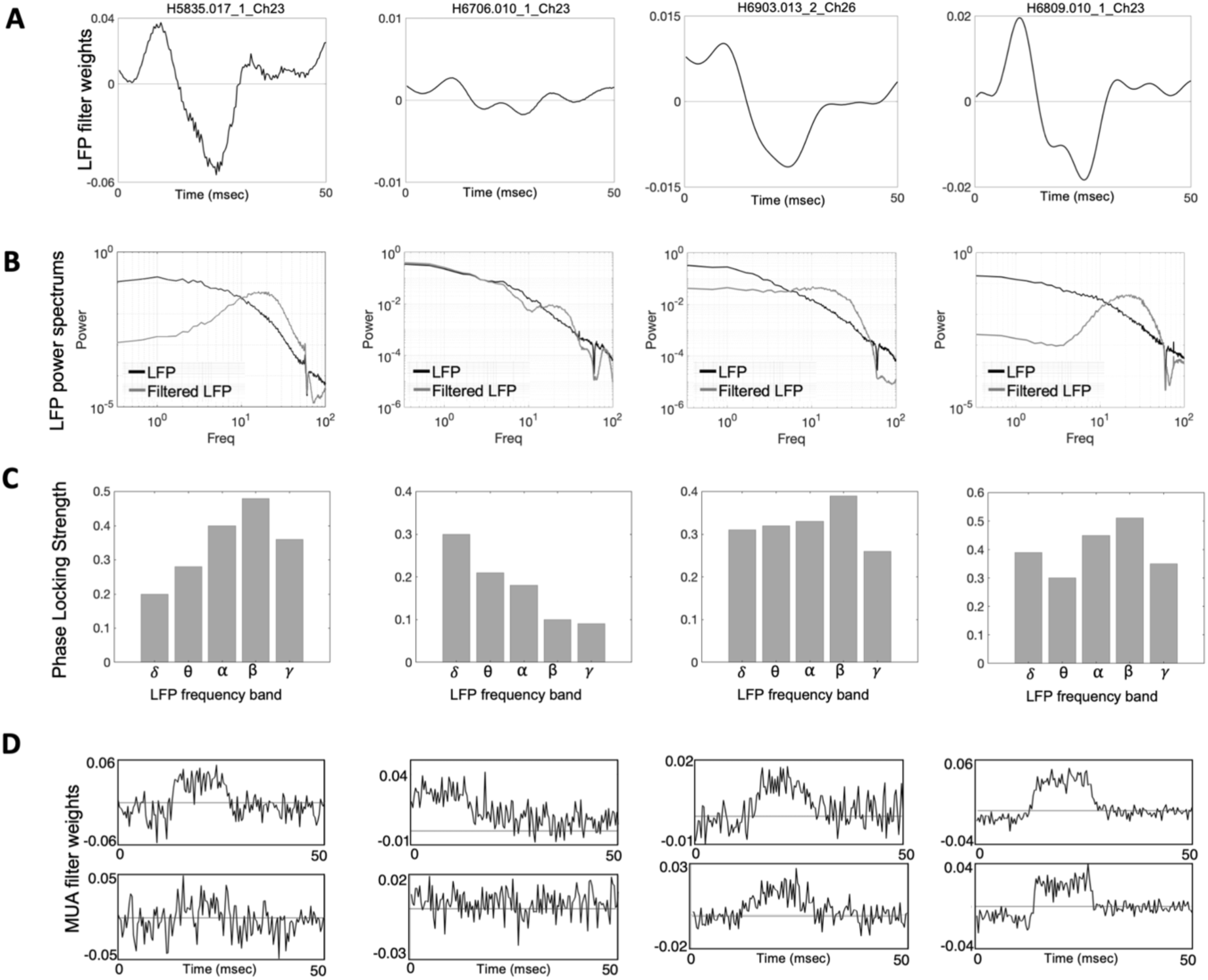
Estimated parameters for cortical state-driven pathways. **A)** Estimated temporal filters from LFP-driven pathway for four example neurons when using LFP from the channel above. **B)** Input LFP (black) and filtered output LFP (grey) from the estimated temporal filters given in panel A. **C)** Phase locking strength (*PLS*) values (comparing spiking activity in relation to the phase of LFP bands). **D)** Estimated temporal filters from MUA-driven pathway for same example neurons when using MUA from the above and below channels.

**S10 Fig.**
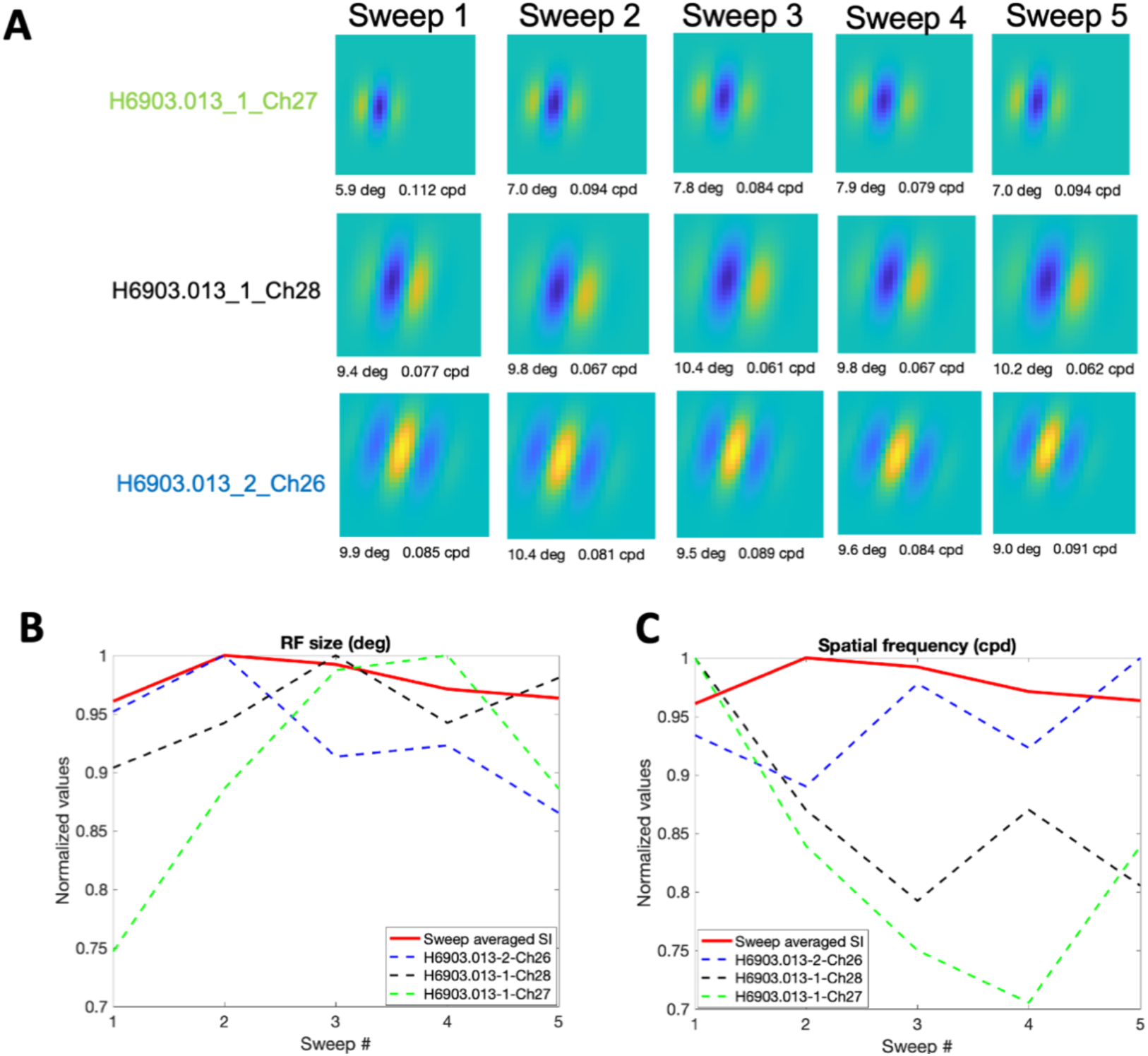
Comparison of RFs with cortical state changes with three example simultaneously recorded neurons. **A)** RF at the peak time lag of three neurons mapped across 5 sweeps. **B)** RF sizes 3 neurons in A across 5 sweeps in dashed lines and cortical state measure SI in red solid line. **C)** Similar to B, the optimal spatial frequencies of 3 neuronal RFs across 5 sweeps vs synchrony index (SI). RF sizes and optimal spatial frequencies were from best-fitting Gabor functions, as described in (Nguyen et al., 2024).

